# A tale of two pumps: Blue light and ABA alter Arabidopsis leaf hydraulics via bundle sheath cells’ H^+^-pumps and channels

**DOI:** 10.1101/2023.03.21.533687

**Authors:** Tanmayee Torne-Srivastava, Yael Grunwald, Mercedes Rosenwald, Ahan Dalal, Adi Yaaran, Veronica Shebtaev, Menachem Moshelion, Nava Moran

## Abstract

This study focuses on the cellular mechanism underlying the co-regulation of the leaf hydraulic conductance (K_leaf_) by blue light (BL) and the stress hormone ABA in *Arabidopsis thaliana*. Our previous work has demonstrated that (1) K_leaf_ increased by BL signaling within the leaf bundle sheath cells (BSCs), which activated their plasmalemma (PM) H^+^-ATPase (AHA2), acidifying the xylem sap; (2) external acidification enhanced the BSCs’ K_leaf_ and their osmotic water permeability (P_f_); (3) ABA decreased both K_leaf_ and P_f_ by reducing the BSCs’ PM aquaporins activity.

We now show, using pH and E_M_ (membrane potential) probes combined with H^+^-pumps inhibitors and manipulations of cytosolic and external Ca^2+^ concentrations ([Ca^2+^]_CYT,_ [Ca^2+^]_EXT,_ respectively), that, in the BSCs: (a) under BL, ABA inhibits AHA2, depolarizing the BSCs and alkalinizing the xylem sap, (b) ABA stimulates the BSCs’ vacuolar H^+^-ATPase (VHA), alkalinizing their cytosol; (c) each pump *stimulation*, AHA2 by BL and VHA by ABA, requires [Ca^2+^]_CYT_ elevation. ABA-effect-mimicking conditions in patch-clamp experiments activate the BSCs’ K^+^-release channels (SKOR and/or GORK). ABA decreased the K_leaf_ of *skor* mutants less than WT’s, while during water deprivation stress, *skor* plants transpired more and their leaves lost relatively less K^+^ than WT. This suggests a role for SKOR in water conservation under drought.

## INTRODUCTION

### The bundle sheath is a regulated barrier to ions and water

Bundle sheath cells (BSCs) enwrap the leaf veins in a tight layer which is increasingly recognized as a selective barrier for ions traveling between the leaf veins and the mesophyll (Shapira et al., 2009; Shapira et al., 2013; Shatil-Cohen and Moshelion, 2012; Wigoda et al., 2017; Niu et al., 2018). BScs constitute also a barrier to water passage between the xylem and the mesophyll, quantifiable as the leaf hydraulic conductance, K_leaf_ (Shatil-Cohen et al., 2011; Grunwald et al., 2021, 2022).

We have shown that K_leaf_ is diminished by ABA applied via the petiole into the xylem of a detached leaf (Shatil-Cohen et al., 2011) and explained the underlying mechanism as an ABA-induced decline of water permeability of the BSCs. We quantified this decline independently in isolated BSCs protoplasts as a diminished P_f_ (osmotic water permeability; Shatil-Cohen et al., 2011), and this was subsequently shown to result from a diminished activity of aquaporins in the BSCs’ plasma membrane (Shatil-Cohen et al., 2011; Sade et al., 2014, Sade et al., 2015; Harayama et al., 2019).

Recently, we reported that K_leaf_ is regulated by the xylem sap pH (pH_EXT_; Grunwald et al., 2021). Whether achieved by pH buffers (ibid.), or by illumination with blue light, which stimulated the BSCs’ H^+^-ATPase, AHA2, via phot receptors (Grunwald et al., 2022), xylem sap acidification increased the K_leaf_, while xylem sap alkalinization (by a pH buffer or by an absence of blue light) decreased it. Similarly, the P_f_ of isolated BSCs’ protoplasts was higher in an acidic solution (pH 6) than in an alkaline solution (pH 7.5; (Grunwald et al., 2021)). Together, these results demonstrated that the BSCs’ external pH (pH_EXT_) affected K_leaf_ by regulating the individual BSCs’ P_f_ (Grunwald et al., 2021). Our finding that pH_EXT_ mediated the *promoting* effect of BL on the BSCs P_f_ and through this – on K_leaf_ (Grunwald et al., 2022), invited a question whether pH_EXT_ mediated also the above-mentioned *inhibitory* effects of ABA on both the BSCs’ P_f_ and K_leaf_ (Shatil-Cohen et al., 2011). Furthermore, since aquaporins are reportedly inhibited by low *cytosolic* pH (Tournaire-Roux et al., 2003) we wondered whether the inhibitory effects of ABA on K_leaf_ involved also the acidification of the BSCs cytosol.

### ABA and apoplast alkalinization

While in response to long-term soil drying the shoot is the source for ABA that appears in the plant, including in the roots (Holbrook et al., 2002; Manzi et al., 2015), w*here* in the leaf is ABA generated in response to water deficit is still debatable (Endo et al., 2008; Bauer, H., Ache, A., Lautner, S., Fromm, J., Hartung, W., Al-Rasheid KA-S., Sonnewald, S ., Sonnewald, U., Kneitz, S., Lachmann, N., Mendel, RR., Bittner, F., Hetherington, AM., and Hedrich, 2013; McAdam, S. A. M. and Brodribb, 2018; Kuromori et al., 2014). Here we focus on resolving *how* ABA – whatever its source – and pH are involved in the *short term* effects on the leaf hydraulics, and therefore we focus on the bundle sheath, a leaf water “valve” in series with stomata which we presume to be regulated independent of stomata (Shatil-Cohen et al., 2011; Grunwald et al., 2021, 2022).

It appears that apoplastic alkalinization is the result of ABA action, rather than the cause of ABA appearance. The reported apoplast alkalinization in a detached leaf petiole-fed with ABA was transient and over within about 10 min - 1 h (in potato, (Hedrich et al., 2001); in *Vicia faba*, (Felle and Hanstein, 2002). The spread of the ABA-induced alkalinization within the leaf apoplast more or less in the vicinity of stomata, was thought to enhance ABA redistribution towards the GCs, but how ABA produced the alkalinization of the GCs apoplast or the leaf apoplast was not conclusively explained (and was even termed “controversial”; (Geilfus et al., 2017). Notwithstanding, current prevalent models of ABA action on GCs include the inhibition of their plasma membrane H^+^-ATPase, AHA1, as the major step towards alkalinization of the guard cell apoplast (Hartung et al., 1988; reviewed by Falhof et al., 2016). As in GCs, ABA applied to root epidermal cells of whole etiolated Arabidopsis seedlings, inhibited the plasma membrane H^+^-ATPase, manifested as an inhibition of H^+^ efflux from seedling roots (Planes et al., 2015).

### Cytosolic pH changes resulting from ABA signaling

Cytosolic pH (pH_cyt_) changes are also a known response to ABA signaling. ABA-induced acidification of the cytosol due to an H^+^-ATPase inhibition was observed in whole Arabidopsis seedlings (Beffagna et al., 1997) and in Arabidopsis epidermal root cells (Planes et al., 2015). In contrast, ABA induced cytosol alkalinization was documented, among others, in corn coleoptiles and parsley hypocotyls and their roots (Gehring et al., 1990), in guard cells (GCs) of orchid (*Paphiopedilum tonsum*) leaf epidermal strips (Irving et al., 1992) and in GCs of *Pisum sativum* (Gonugunta et al., 2008), of Arabidopsis (Bak et al., 2013), and of *Vicia faba* (Blatt and Armstrong, 1993). This ABA-induced cytosol alkalinization was attributed to the ABA-induced stimulation of the activity of the GCs’ vacuolar V-type H^+^-ATPase, VHA (Bak et al., 2013), the underlying mechanism of which is still not completely resolved (Jezek and Blatt, 2017).

### ABA and blue light signaling pathways in the BSCs

We chose the GC model to guide our exploration of signaling in the BSCs, since a GC is also a leaf cell and therefore perhaps closer to a BSC than a root cell or a pollen tube, but especially because GCs have been the most-extensively studied leaf cell species. This choice has been supported also by our recent discovery of an autonomous blue light (BL) signaling pathway in the BSCs, resembling that in the GCs (Grunwald et al., 2022). In the BSCs, as in GCs, the perception of BL by phot1 and phot2 with the participation of BLUS1, led to the activation of a plasma membrane H^+^-ATPase (AHA2 in the BSCs). The activation of the BScs’ AHA2 led further to xylem sap acidification, cytosol alkalinization, the increase of the BSCs’ P_f_ and of K_leaf_. Here, we extend our recent model of BL signaling within the BSCs (ibid.) beyond that of GCs (Inoue and Kinoshita, 2017) by testing a *hypothesis* that the BL-induced activation of the BSCs’ AHA2 depends on elevated cytosolic Ca^2+^ concentration ([Ca^2+^]_CYT_). The rationale for this stems from reports (reviewed by Harada and Shimazaki, 2007) of, among others, BL increase of [Ca^2+^]_CYT_ mediated by phot1 receptors in Arabidopsis seedlings (Baum et al., 1999) or by phot1 and phot2 in Arabidopsis rosettes (Harada et al., 2003), and even more specifically, on phot-mediated BL activation of Ca^2+^ channels in Arabidopsis mesophyll cells (Stoelzle et al., 2003) or BL activation of Ca^2+^ inward fluxes mediated by phot1 in Arabidopsis seedlings (Babourina et al., 2002).

The ABA signaling cascade leading to the closure of stomata, is initiated in the GCs by ABA binding to its intracellular receptor(s) (PYR/PYL), recruiting a clade A protein phosphatase PP2C (e.g., ABI1) to the complex and relieving the inhibition of kinases downstream (Ma et al., 2009; Umezawa et al., 2009). A number of steps downstream, the GCs alkalinize their apoplast (Geilfus, 2017) and their cytosol (Blatt and Armstrong, 1993). In *abi1-1* plants (Koornneef et al., 1984; Meyer et al., 1994), the mutated PP2C protein (*abi1-1*) cannot be recruited to the ABA/PYR/PYL complex upon ABA binding (due to an amino acid substitution G180D, (Leung, J., Merlot, S., and Giraudat, 1997), but still retains its phosphatase activity (Merlot et al., 2001; Imes, D., Mumm, P., Böhm, J., Al-Rasheid, K. A.S., Marten, I., Geiger, D., and Hedrich, 2013). The mutation is thus dominant-negative: it enables the circulation of an active protein phosphatase allowing it to continue its negative regulation of the downstream ABA signaling pathway (Gosti et al., 1999; Cai et al., 2017, and references therein), rendering the guard cells (among other cells) ABA insensitive (Wu et al., 2003) which eventually prevents the GC’s AHA1 inhibition by ABA and averts the GCs depolarization.

### Leaf hydraulic conductance and K^+^ channels

ABA signaling in the GCs may also include the rise of cytosolic Ca^2+^ concentration ([Ca^2+^]_CYT_; McAinsh et al., 1990) and it includes the opening of anion channels (e.g.,Levchenko et al., 2005) and of the GCs’ sole K^+^-release (K_OUT_) channel GORK (Guard cell Outward Rectifying K^+^ channel; Ache et al., 2000; Hosy et al., 2003), leading to the efflux of anions and K^+^. Thus, we test here an additional *hypothesis*, guided by the GCs model, that the water-deficit-signaling ABA acting on the the Arabidopsis leaf BSCs, not only alkalinizes their apoplast, i.e., the leaf xylem sap, and their cytosol, as well as depolarizes its membrane (due to lower AHA activity) but that both the pH and depolarization signals enhance the activity of the BSCs’ K_OUT_ channels leading to K^+^ efflux into the BSCs’ apoplast (as in GCs), possibly increasing the [K^+^] in the leaf veins, at the expense of [K^+^] in the surrounding cells. Notably, the BSCs harbor both, GORK and SKOR (Stelar K^+^ Outward Rectifier, the other Arabidopsis K_OUT_ channel; Gaymard et al., 1998), and both are expressed also in the mesophyll (beside several other K^+^ channels; Wigoda et al., 2017). Furthermore, we *hypothesize* that ABA signaling in the BSCs of *skor*, a SKOR mutant plant, releases less K^+^ into the leaf veins, compared to WT.

Overall, our experiments confirm our general hypothesis that BSCs, like GCs, possess autonomous BL and ABA signalling cascades, wherein BL upregulates the AHA2 and ABA regulates two BSCs’ proton pumps. Beside the similarities we find also a few differences between the GCs and the BSCs.

Taken together, our findings are *consistent* (1) with an *up*regulation by BL of a leaf “hydraulic valve” other than guard cells – the BSCs layer – and (2) with an *down*regulation of this “hydraulic valve” by ABA. We propose that the first occurs by the *up*regulation of the BSCs’ aquaporin activity and the second, by the *down*regulation of the BSCs’ aquaporin activity and *up*regulation of their SKOR activity. By retaining the xylem water, the downregulation of the hydraulic valve might help avert drought-stress-induced embolism.

## RESULTS

### Light-enhanced leaf hydraulic conductance depends on free Ca^2+^ concentration in the BSCs’ cytosol

Since BL increased K_leaf_, and since it also increased the free Ca^2+^ concentration in the cytosol ([Ca^2+^]_CYT_) of various Arabidopsis cells, we asked whether this K_leaf_ increase depended on an elevated [Ca^2+^]_CYT_ in the BSCs. When the membrane-permeant BAPTA-AM (a Ca^2+^-chelator activated within the cytosol) was added to the artificial xylem sap (AXS) perfused via the petiole of a detached and illuminated WT Arabidopsis (Col) leaf, its K_leaf_ decreased considerably (Fig. 1A, Suppl. Fig. S1). This suggested that an undiminished (and very likely – BL-elevated) [Ca^2+^]_CYT_ level in the cells in the xylem vicinity – presumably, in the BSCs – is important for attaining a high K_leaf_.

**FIGURE 1.**
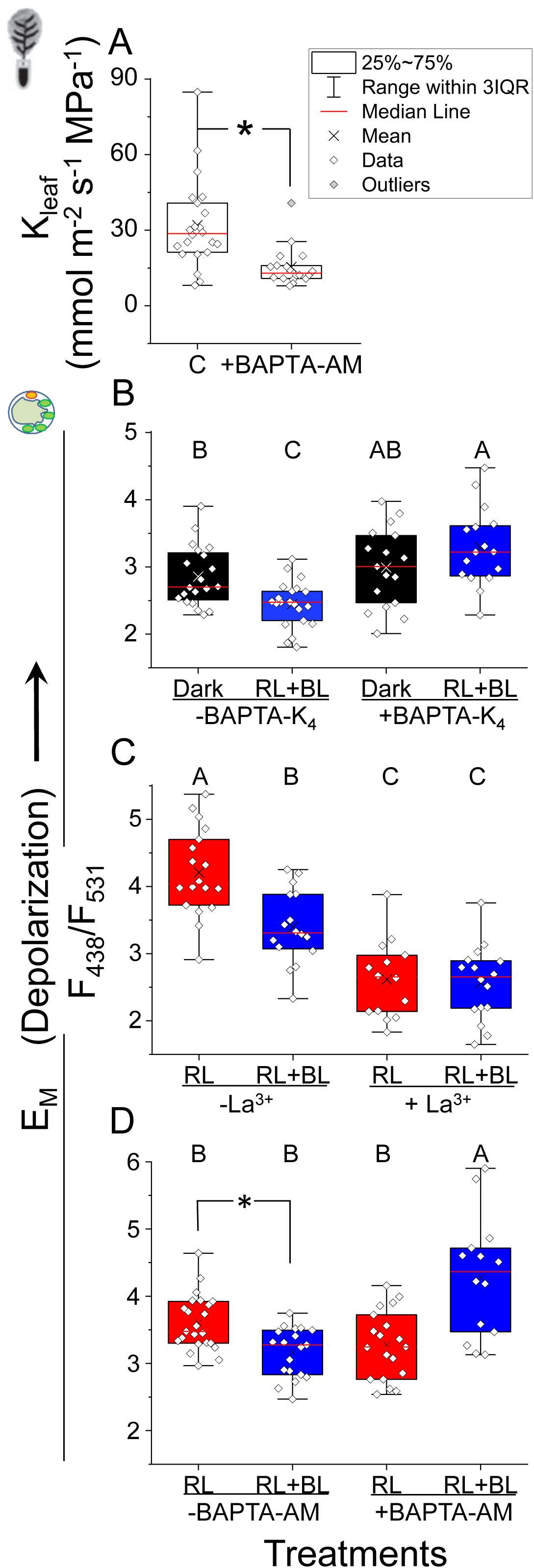
The hydraulic conductance (K_leaf_) of a detached leaf and the underlying BL-activation of AHA2, both depend on BL-elevated cytosolic free Ca^2+^ concentration ([Ca^2+^]_CYT_). A. BAPTA-AM (10 μM), the cytosolic Ca^2+^ chelator, administered in an artificial xylem solution (AXS) for 1.5-2 hs to a detached leaf of Arabidopsis WT (Col) via the petiole, diminishes the K_leaf_ relative to control (C). See the data used for K_leaf_ calculation in Suppl. Fig. S1. *: a difference from control (C; P = 0.000291, by a two-tailed t-test, equal variance; outliers excluded). **Inset**, detached leaf schematics. **B-D**. Manipulations intended to decrease [Ca^2+^]_CYT_ abolish the blue-light (BL)-induced hyperpolarization (i.e., the BL-induced decrease of membrane potential, E_M_) of WT (Col) BSCs protoplasts (biological repeats). E_M_ was determined using the fluorescent dual-excitation ratiometric dye, di-8-ANEPPS (30 μM). See also the Suppl. Fig. S2. **B.** The membrane-*im*permeant (ionic) Ca^2+^ chelator BAPTA-K_4_ (1 mM), which lowered the free [Ca^2+^] in the bath from 0.4 mM down to approx. 0.6 μM, abolished the red+blue light (RL+BL)-induced hyperpolarization. Different letters indicate statistically different means (ANOVA, Tukey-HSD test, P<0.05). **Inset**: an isolated BSC protoplast schematics. **C**. The BL-induced hyperpolarization was abolished by the Ca^2+^ channel blocker, La^3+^ (50 μM). **D**. BAPTA-AM (10 μM) also abolished the BL-induced hyperpolarization. *: a significant difference (P= 0.000888) by a t-test as above.

### Ca^2+^ influx is a prerequisite for the blue light activation of the BSCs’ AHA2

To test whether BL activation of the BSCs’ AHA2 involves Ca^2+^ influx into the cytosol and elevation of [Ca^2+^]_CYT_, we monitored the BSCs’ AHA proton-extrusion activity by monitoring the membrane potential (E_M_) of BSCs protoplasts using an E_M_ fluorescent probe, di-8-ANEPPS without or with three treatments aimed at antagonizing the BL-induced potential elevation of [Ca^2+^]_CYT_: (a) direct chelation of the external Ca^2+^ by the bath-applied ionic BAPTA-K_4_, which lowered the external Ca^2+^ concentration ([Ca^2+^]_EXT_) by three orders of magnitude from 0.4 mM to approx. 0.6 μM, or (b) adding the Ca^2+^ channel blocker, La^3+^, or (c) the chelation of cytosolic Ca^2+^ by BAPTA-AM. While in the controls, BL evoked hyperpolarization, compared to either dark or to RL alone (Figs. 1B-1D, left), each of these treatments abolished it (Figs. 1B-1D, right; Incidentally, RL and darkness did not differ with regard to AHA2 activity and BL hyperpolarized the BSCs and alkalinized their cytosol relative to both RL and dark (see Fig. 5A and 5B in (Grunwald et al., 2022). Together, these results suggest that Ca^2+^ influx via Ca^2+^ channels is required for the BL-induced stimulation of the BSCs’ H^+^-ATPase, AHA2, previously established as responsible for the BL-induced BSCs hyperpolarization.

### ABA alkalinizes the xylem perfusate due to AHA2 inhibition

To test our hypothesis that ABA inhibits the BSCs’ AHA2, we assayed the BSCs responses to ABA under the same illumination (RL+BL) as before. We monitored the pH within the xylem (pH_EXT_) of detached illuminated leaves as reflecting the AHA proton-extrusion activity of the BSCs. The pH_EXT_ of WT (Col) leaves, perfused with a xylem perfusion solution (XPS) without ABA was approx. 5.8 (Fig. 2A). Inclusion of 10 μM ABA in the XPS, elevated the mean pH_EXT_ by about 0.4 pH units (from pH 5.8 to 6.2; Fig. 2A), suggesting that ABA inhibited net proton extrusion from the BSCs into the xylem. In contrast, ABA did not affect the pH_EXT_ in the leaves of *aha2-4* (Col), a knockdown mutant of AHA2 (Haruta et al., 2010, 2012; it remained in the range of 6.0 to 5.7; Fig. 2B), suggesting that in the absence of AHA2 activity, ABA did not affect proton extrusion via the plasma membrane towards the xylem. Moreover, in *aha2-4* complemented with *SCR:AHA2*, with AHA2 active exclusively in the BSCs (Grunwald et al., 2021), ABA treatment alkalized pH_EXT_ by 0.8 units (from 5.6 to 6.4; Fig. 2C), doubly that in WT, in support of our hypothesis that in the BSCs, AHA2 is the target of ABA-induced inhibition.

**FIGURE 2.**
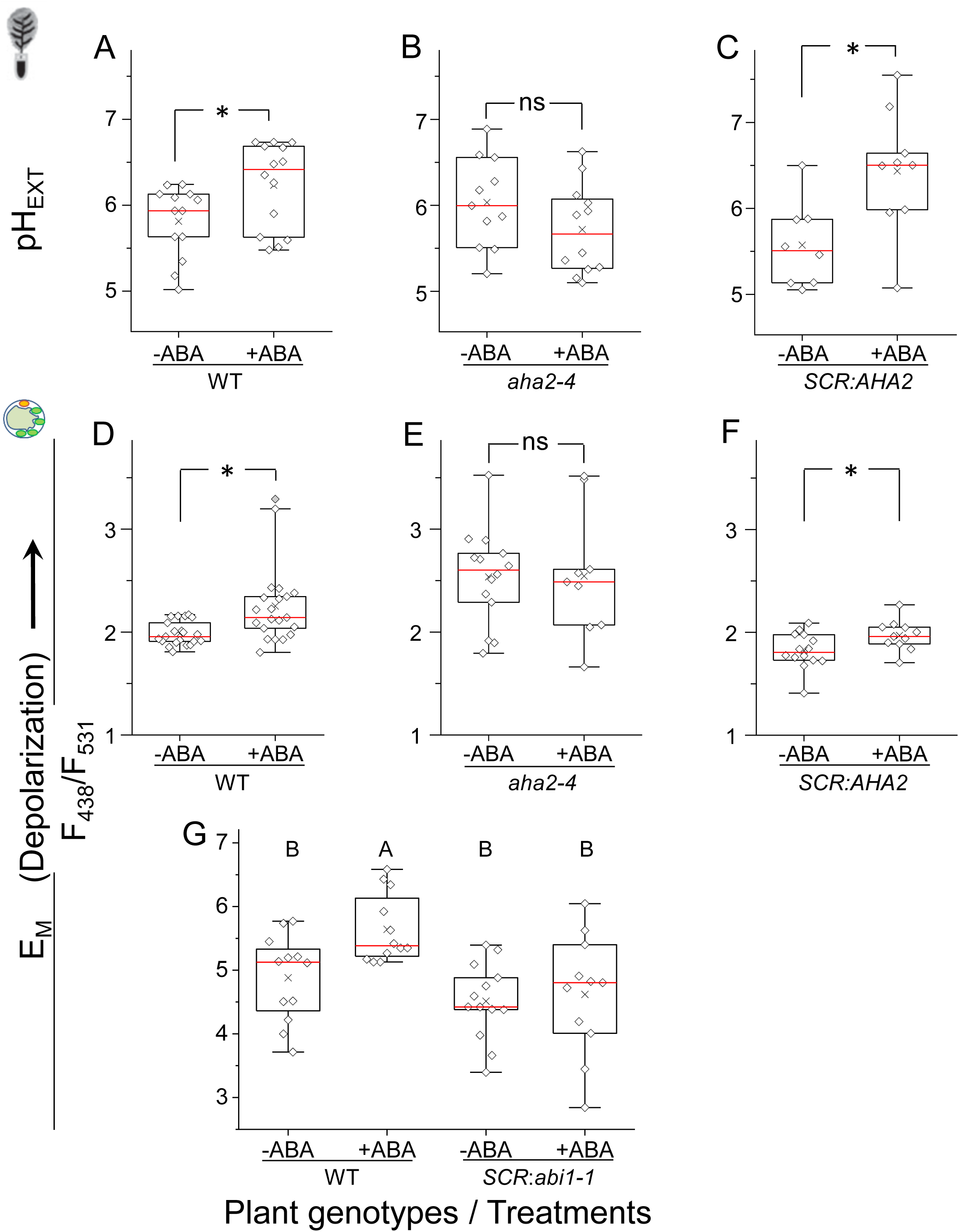
ABA alkalinization of xylem perfusate and depolarization of BSCs are due to AHA2 inhibition by an ABA signaling path involving the early element, ABI1. (*ABA-insensitive 1*, a protein phosphatase 2C, PP2C). A-C. ABA (10 μM ) elevates the pH of the xylem perfusion solution, XPS (pH_EXT_) in the detached leaves of Arabidopsis plants only in the presence of active AHA2 in the BSCs. pH_EXT_ was determined using the fluorescent dual-excitation ratiometric dye, FITC-D (10 μM). A. WT (Col). *: a significant difference (P=0.02596). B. *aha2-4* (Col), a line of a knockdown mutant of the AHA2. ns: a non-significant difference. C. *SCR:AHA2*: *aha2-4* mutant complemented with *SCR:AHA2* under the BSCs-directing promoter *SCR* (*Scarecrow*). *: a significant difference (P=0.01238). D-F. ABA (3 μM) depolarizes the BSCs (elevates their membrane potential, E_M_). D. WT (Col) BSCs protoplasts. *: a significant ABA-induced depolarization (P=0. 0.00412). Inset: a RL+BL-illuminated BSC protoplast schematics. E. *aha2-4* mutant. ns: not significant. F. *SCR:AHA2* plants as in C. *: a significant ABA-induced depolarization (P=0.03311). G. ABI1 mediates the ABA-induced BSCs’ AHA2 inhibition. ABA (3 μM) depolarizes BSCs protoplasts isolated from the leaves of WT (Col) Arabidopsis but fails to change the E_M_ of BSCs from transgenic WT (Col) plants transformed with *SCR:abi1-1*, the mutated *ABI1-1* (*ABA Insensitive1-1)* gene. See also Suppl. Fig. S3.

### ABA-induced BSCs depolarization results from AHA2 inhibition

To test the *prediction* that ABA inhibition of the BSCs’ will cause their depolarization, we applied 3 μM ABA to WT (Col) BSCs protoplasts under RL+BL illumination and monitored their membrane potential, E_M_ (as in Fig. 1). As expected, ABA depolarized the WT (Col) BSCs protoplasts (Fig. 2D). In contrast, ABA did not depolarize the BSCs protoplasts of *aha2-4* (Col) which were *a priori* practically devoid of AHA2 activity (Fig. 2E), whereas, in *aha2-4* complemented with *SCR:AHA2*, with the missing AHA2 activity restored exclusively to the BSCs, ABA-induced BSCs depolarization was also restored (Fig. 2F). Together, these results suggest that the depolarization is due specifically to AHA2 inhibition by ABA (and that the activity of AHA2, specifically, is required as a background in order to demonstrate this inhibition). Interestingly, the depolarization was evident during minutes 10-15 after the BSCs exposure to ABA, but not during the first 5 min (Suppl. Figs. S2A, S2B).

### ABA-induced AHA2 inhibition involves the protein phosphatase ABI1

Since the protein-phosphatase 2C, ABI1, mediates the classical ABA signaling sequence in guard cells, we examined its role in the ABA-signaling in the BSCs. We compared the ABA effect on the membrane potential of single BSCs protoplasts isolated from WT (Col) plants to that on BSCs protoplasts from transgenic *SCR: abi1-1* WT (Col) plants i.e., plants transformed with a mutated ABI1 gene under the BSCs-directing *SCR* promoter. The ABA-induced depolarization observed in the WT BSCs was abolished in the *SCR:abi1-1* BSCs (Fig. 2G), demonstrating the autonomy of the BSCS’s ABA-signaling pathway which inhibits AHA2. In support of these results, ABA failed to elicit XPS alkalinization in the detached leaves of *abi1-1* (Ler) mutant plants (Suppl. Figs. S3A, 3B) and the transgenic *SCR-abi1-1* (Col) plants (Suppl. Fig. S3C, 3D).

### ABA alkalinizes the cytosol in spite of AHA2 inhibition

We showed recently that the blue light-activated AHA2 alkalinized the BSCs’ cytosol. Therefore, by extension, we expected the inverse, i.e., a cytosolic acidification, to result from AHA2 inhibition by ABA. To test this, we applied ABA and monitored the cytosolic pH (pH_CYT_) using the ratiometric, dual-emission fluorescent pH probe, SNARF1 (Suppl. Figs. S4A-S4C). Contrary to this simple prediction, ABA (3 μM) added in the bath, did not acidify, but rather, alkalinized the cytosol of isolated WT (Col) BSCs protoplasts within 15 minutes by 0.9 pH units (from 7.7 to 8.6, Fig. 3A, left). This suggests that the BSCs’ cytosol alkalinization does not depend on their AHA2 inhibition. Importantly, the ABA-induced alkalinization was well advanced already within the first 5 minutes after ABA introduction to the bath (Suppl. Fig. S4D) and persisted at least until min 15 after ABA exposure (Suppl. Fig. S4E).

**FIGURE 3.**
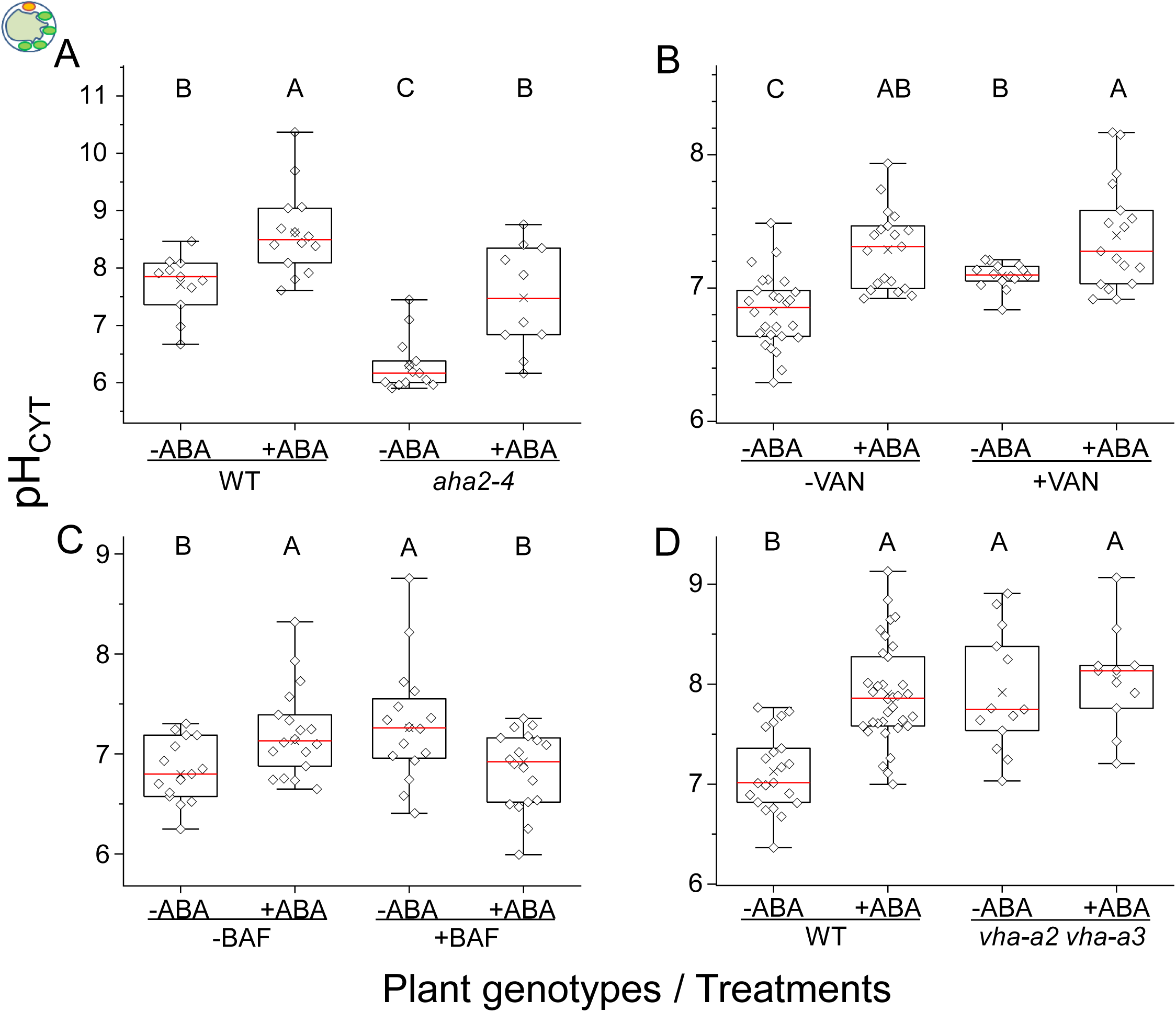
ABA-induced cytosol alkalinization does not involve AHA2, but is mediated via VHA. Cytosol pH (pH_CYT_) of BSCs protoplasts was determined using SNARF1 (Suppl. Fig. S4). **A.** *AHA2* knockdown has no effect on ABA (3 μM)-induced increase of pH_CYT_ in BSCs protoplasts from WT (Col) or from *aha2-4* (Col) mutant plants ABA. The F-ratio values were converted to pH_CYT_ using the calibration curve of Suppl. Fig. S4C. **Inset**: a RL+BL-illuminated BSC protoplast schematics. **B.** The AHA inhibitor, vanadate (VAN, 1 mM), has no effect on the ABA response in WT (Col) BSC protoplasts. The F-ratio values were converted to pH_CYT_ using the calibration curve of Suppl. Fig. S4B. **C.** The vacuolar H^+^-ATPase (VHA)-specific inhibitor, bafilomycin A1 (BAF, 100 nM), abolishes the ABA response in WT (Col) protoplasts. Note the pH_CYT_ increase under ABA alone relative to control (-ABA,-BAF) and its abolishment under ABA and BAF together. **D.** The double VHA mutation *vha-a2 vha-a3* (Col) abolishes the pH_CYT_ response to ABA.

In accord with the above prediction, the “resting” pH_CYT_ of the *aha2-4* mutant was indeed considerably lower than the WT’s pH_CYT_ , by 1.4 pH units (the mutant’s 6.3 vs WT’s 7.7; Fig. 3A). Nevertheless, treating the *aha2-4* mutant BSCs with ABA replicated the effect found in WT, also causing a pronounced alkalinization, by 1.2 pH units (roughly from 6.3 to 7.5, Fig. 3A, right), definitively ruling out any involvement of AHA2 in the ABA-induced BSCs’ cytosol alkalinization.

To narrow down the candidacy of other proton pumps potentially responsible for this fast ABA-induced cytosol alkalinization, we pre-incubated the WT (Col) BSCs protoplasts with 1 mM vanadate (a known general P-type ATPases inhibitor). Surprisingly, when we applied vanadate by itself to WT (Col) BSCs, to mimic the *aha2-4* mutation, it elevated the pH_CYT_ relative to control (C) BSCs (from 6.8 to 7.1; Fig. 3B) which contrasted sharply with the abovementioned pH_CYT_-lowering effect of knocking down AHA2 in the *aha2-4* mutant. However, similarly to the *aha2-4* mutation, vanadate did not prevent, the *ABA-induced increase* of the BSCs pH_CYT_. When ABA was added to the vanadate-treated WT protoplasts, pH_CYT_ increased by approximately 0.3 pH units relative to vanadate alone (from about 7.1 to 7.4, Fig. 3B, right). This cytosol alkalinization by ABA in the vanadate background ruled out the involvement of any vanadate-susceptible P-type H^+^-ATPases (including the AHA2) (Morsomme and Boutry, 2000; Palmgren, 2001) in the ABA-induced BSCs’ cytosol alkalinization, implicating instead a vanadate-*in*sensitive mechanism extruding protons from the cytosol, such as the vacuolar H^+^-ATPase, VHA, and/ or the vacuolar pyrophosphatase (PP-ase), APV1(Krebs, M., Beyhl, D., Görlich, E., Al-Rasheid, K. A. S., Marten, I., Stierhof, Y.D., Hedrich, R., and Schumacher, 2010).

### The ABA-induced cytosol alkalinization is mediated by VHA

To explore directly the involvement of VHA in the ABA-induced cytosol alkalinization under BL, we used the VHA-specific inhibitor, bafilomycin A1. Interestingly, 100 nM bafilomycin A1 added to the bath without ABA, elevated the pH_CYT_ within minutes, by ∼0.4 pH units relative to the control (from about 7.2 to 7.6, Fig. 3C), not much different from the WT’s response to ABA (from 7.2 to 7.5, Fig. 3C). However, when bafilomycin A1 was added to the bath together with ABA, pH_CYT_ rise was abolished (its value remained approx. 7.2, Fig. 3C). Surprisingly, pH_CYT_ did not decline below the pH_CYT_ value in the control, as could be expected, given that both VHA and AHA2 were inhibited simultaneously. In a control experiment, we eliminated any interference of the solvent, DMSO in the results (Suppl. Fig. S5).

In a complementary experiment, we assayed the effect of ABA on the double mutant of the two tonoplast-located isoforms of the a subunit of VHA (*vha-a2 vha-a3* (Col); Krebs et al., 2010). The “resting” pH_CYT_ of the BSCs in this VHA-activity-devoid mutant was more alkaline than in WT (Col), by about 0.7 pH units (roughly, 7.9 vs. 7.2, respectively; Fig. 3D). This may be attributed to the life-long disruption of homeostasis of the vacuolar pH, pH_VAC_, likely compensated by AHA activity and possibly also by the activity of VHA-a1 in the trans-Golgi network/early endosome (TGN/EE)-derived vesicles (which could be also responsible for the viability of the double mutant; (Krebs, M., Beyhl, D., Görlich, E., Al-Rasheid, K. A. S., Marten, I., Stierhof, Y.D., Hedrich, R., and Schumacher, 2010). ABA treatment of the double-mutant did not affect its pH_CYT_ (with pH_CYT_ remaining roughly at 7.9-8.1, Fig. 3D), suggesting, again, that the fast, ABA-induced elevation of pH_CYT_ in the WT requires an active tonoplast VHA-a2 VHA-a3.

### ABA-induced cytosol alkalinization is not a prerequisite for AHA2 inhibition

To resolve whether AHA2 inhibition by ABA depends on cytosol alkalinization (due to a concomitant stimulation of VHA), we monitored E_M_ with di-8-ANEPPS without and with added ABA, in BSCs treated with bafilomycin A1 to inhibit VHA activity. In contrast to the above-mentioned fast cytosolic alkalinization induced by ABA (Suppl. Fig. S4), no change of E_M_ could be seen during the first 5 minutes after ABA was flushed into the bath (Suppl. Fig. S2C). At minutes 10-15, however, the depolarization was already apparent, indicating the inhibition of AHA2 in spite of the absence of VHA activity (Suppl. Fig. S2D).

### The BSCs’ VHA regulation

#### Neither light (RL or BL), nor decline of [Ca^2+^]_CYT_ affect pH_CYT_

To find out whether the BSCs’ VHA is activated by BL in a Ca^2+^-dependent manner, similarly to AHA2, we treated WT (Col) BSCs protoplasts by vanadate (1 mM) – to eliminate a possible effect of AHAs on pH_CYT_ – and monitored their pH_CYT_ under RL without or with BL, without and with BAPTA-AM (Fig. 4A). We found no difference between the treatments (pH_CYT_ remained in the range of 7.8-8.2), suggesting that VHA is not affected by the BL-induced [Ca^2+^]_CYT_ increase, nor is it sensitive to a *decrease* of [Ca^2+^]_CYT_ by chelation below the “resting” level.

**FIGURE 4.**
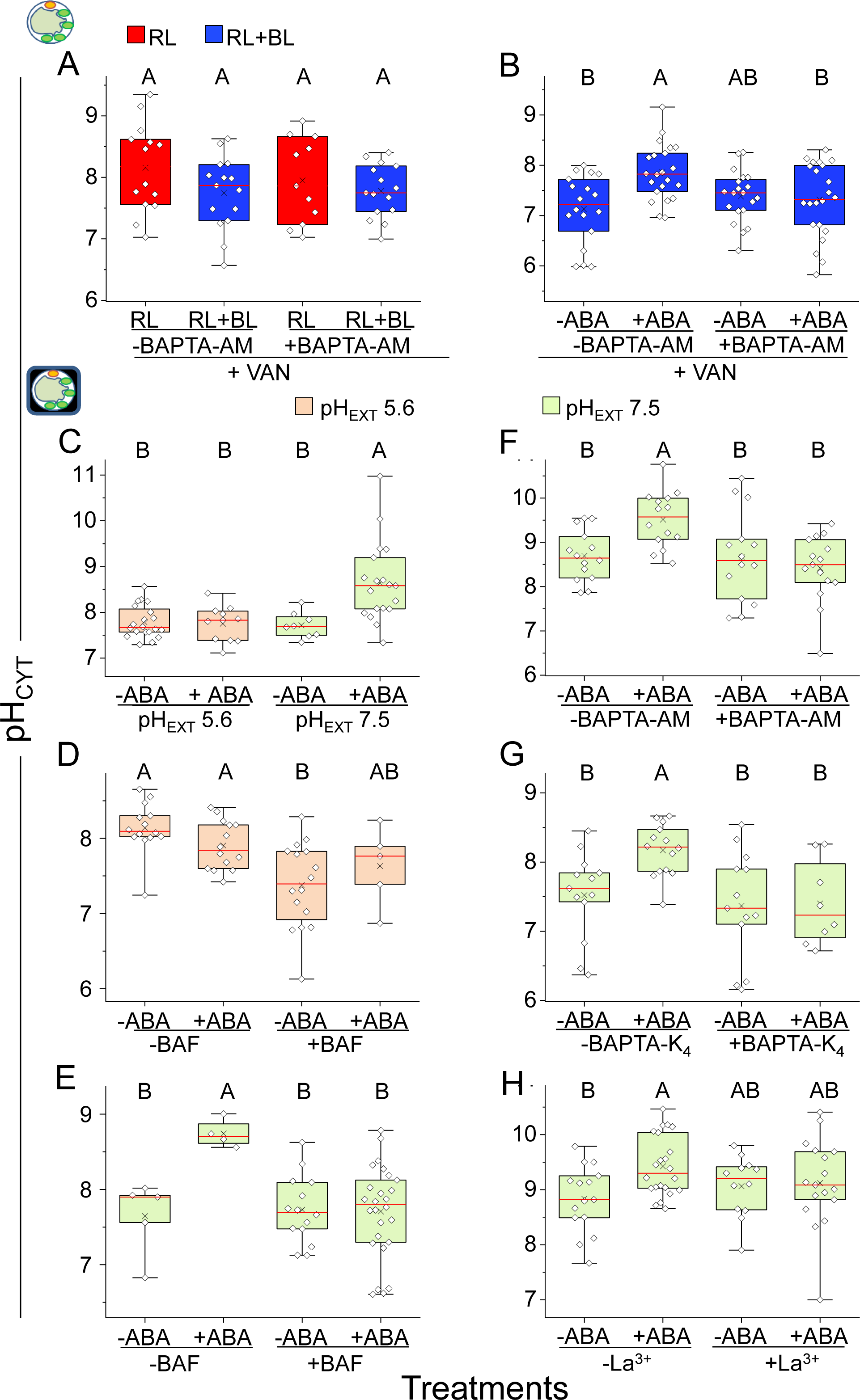
Neither light (RL or BL), nor [Ca^2+^]_CYT_ decline affect VHA activity, but ABA-induced [Ca^2+^]_CYT_ rise – does, requiring light or alkaline pH_EXT_. A-B. In the presence of the AHA inhibitor, vanadate. A. BL has no effect on pH_CYT_ in WT (Col) BSCs protoplasts and [Ca^2+^]_CYT_ chelation by BAPTA-AM does not alter pH_CYT_ irrespective of illumination. Inset. An illuminated BSC protoplast schematics. B. BAPTA-AM abolishes the ABA-stimulated cytosol alkalinization in WT (Ler) BSCs protoplasts under RL+BL. C-H. In the AHA2-inactivating darkness. Inset: a protoplast-in-the-dark schematics. C. ABA (3 µM) elevates pH_CYT_ in WT (Ler) BSCs protoplasts only at pH_EXT_ 7.5, not at pH_EXT_ 5.6. D. Bafilomycin A1 (BAF, 100 nM), applied by itself at pH_EXT_ 5.6 lowers the pH_CYT_, suggesting a basal activity of VHA at pH_EXT_ 5.6, in the dark. E. At pH_EXT_ 7.5, ABA-induced cytosol alkalinization is abolished by bafilomycin A1, consistent with VHA stimulation by ABA, while bafilomycin A1 by itself has no effect on pH_CYT_. F. At pH_EXT_ 7.5, in an otherwise standard bath solution, the [Ca^2+^]_CYT_ chelator, BAPTA-AM (10 μM) abolishes the pH_CYT_ increase by ABA in WT (Ler) BSCs protoplasts. G. The ABA-induced pH_CYT_ elevation is abolished in WT (Ler) BSCs protoplasts by decreasing [Ca^2+^]_EXT_ in the bath solution (at pH_EXT_ 7.5 and modified as in Fig. 1B) by the external, ionic Ca^2+^ chelator, BAPTA-K_4_ (1 mM; the calculated free [Ca^2+^]_EXT_ was approx. 0.04 μM). H. The ABA-induced pH_CYT_ rise is abolished in WT (Col) BSCs by the Ca^2+^ channel blocker, La^3+^ in the bath solution (at pH_EXT_ 7.5 and modified as in Fig. 1C).

**Figure 5.**
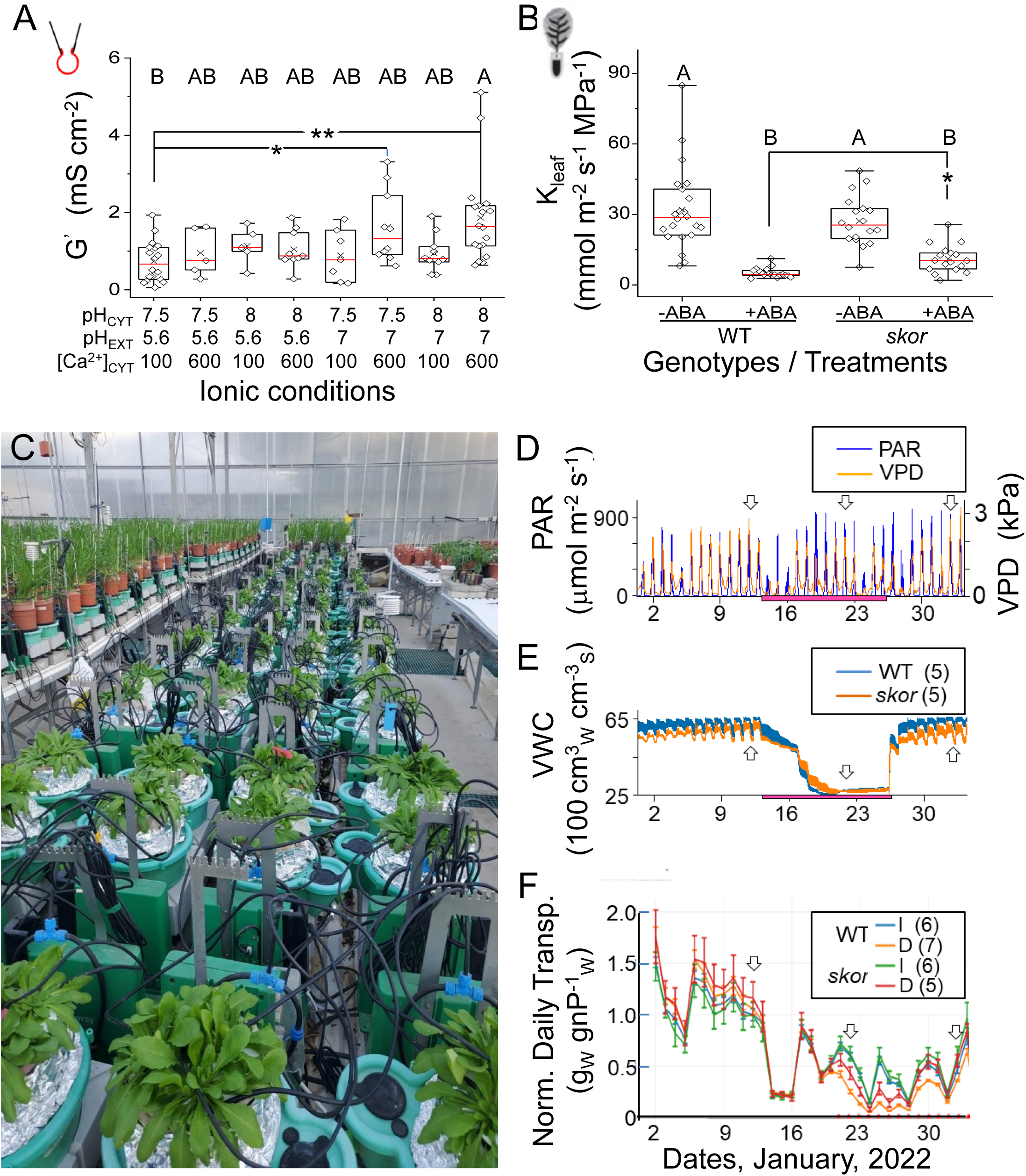
The K^+^-release (K_OUT_) channel SKOR is involved in the regulation of transpiration under stress. **A.** K^+^-release channel activity in patch-clamped WT (Ler) BSCs protoplasts is represented by an ensemble of specific membrane conductances (G’) determined at an extreme depolarizing membrane potential (E_M_ = +117 or +97 mV), is shown as a function of the ionic milieu near the membrane (attained by various combinations of bath pH (pH_EXT_), pipette pH (pH_CYT_) and pipette free Ca^2+^ conc. ([Ca^2+^]_CYT_)). Different letters indicate significantly different G’ means (ANOVA, as in previous figures). Asterisks mark significantly different means by t-test (*: P=3.4E-3, **: P= 9.4E-4). Note that the most enhanced G’ of the K^+^-release channels was attained only at the non-acidic external pH and the high [Ca^2+^]_CYT_ (pH_EXT_ 7, pH_CYT_ 7.5 or 8 and [Ca^2+^]_CYT_ 600 nM). **Inset**. The whole-cell recording configuration schematics. **B.** The effect of ABA on the hydraulic conductance (K_leaf_) of detached leaves. *: a significant difference at P = 0.000565 (by t-test as before). Note the attenuated decline of K_leaf_ in the *skor* (Col) mutant relative to WT (Col). See also the Suppl. Fig **S**6. **C.** A photo of a physiology-phenotyping greenhouse for whole Arabidopsis plants (seen in the central table; winter 2022). The plants were grown in large pots (4 plants ea.) and their growth and daily transpiration were monitored continuously in a greenhouse using lysimeters and additional sensors. **D-F.** 33-days-long simultaneous records from the phenotyping greenhouse (2 Jan – 3 Feb, 2022). **D.** Atmospheric conditions, recorded by the weather station: PAR (photosynthetically active radiation) light, VPD (vapor pressure difference). **E.** The mean (± SE) volumetric water content (VWC), i.e., the volume of water (cm^3^_W_) contained in the drying soil relative to the initial volume (soil + water) of the maximally wet soil (cm^3^_S_), derived from the reported measurements by soil sensors from the pots of both droughted genotypes. Note the decline in VWC during the withdrawal of irrigation indicated by the magenta horizontal bar; the last irrigation occurred in the night between the 13^th^ and the 14^th^ of Jan 2022, and irrigation was resumed in the night between the 26^th^ and the 27^th^ of Jan 2022, followed by 8 days of recovery. **F.** The mean (± SE) normalized daily transpiration (Norm. Daily Transp.) of continuously irrigated (I) and water-deprived (droughted, D) WT (Col) plants and *skor* (Col) mutant plants. The Norm. Daily Transp. values (biological repeats) were derived from the combined daily transpiration of the four plants in a pot (the weight of lost water, g_W_) normalized to their corresponding calculated combined net weight (g_nWP_). Red dots indicate a significant difference (P<0.05, by t-test) between the means of the *droughted* plants of the two genotypes, observable first on 21^st^ Jan, the 8^th^ day of drought. Note the slightly higher normalized daily transpiration of the *droughted skor* mutant plants relative to the *droughted* WT plants during the late phase of the drought period (in spite of the identity of their VWC), lasting into the recovery period. Small vertical arrows indicate representative dates the data of which are shown expanded in Suppl. Fig. S7.

#### In the light, VHA response to ABA requires elevated [Ca^2+^]_CYT_

To resolve whether VHA stimulation by ABA depended on [Ca^2+^]_CYT_ (as suggested by Tang et al., 2012), see further down in the Discussion), we assessed the effect of BAPTA-AM on the ABA response in WT (Ler) BSCs protoplasts illuminated by RL+BL, again, using vanadate (1 mM) to eliminate a potential intereference of AHA. As expected, ABA alkalinized the cytosol (by 0.6 pH unit, from pH 7.2 to 7.8; Fig. 4B, left). BAPTA-AM, however, abolished this ABA-induced alkalinization (pH_CYT_ remained around 7.4, Fig. 4B, right), suggesting that at least one of the signaling steps in the VHA stimulation by ABA depends on the elevation of [Ca^2+^]_CYT_.

#### In the dark, VHA response to ABA requires alkaline pH_EXT_ and elevated [Ca^2+^]_CYT_

Using darkness as an alternative means to eliminate the activity of AHA (Grunwald et al., 2022), we examined the effect of ABA on pH_CYT_ in darkness to test whether the BSCs’ VHA required illumination in order to be stimulated by ABA. ABA did not alkalinize the BSCs’ pH_CYT_ (which remained around 7.7, Fig. 4C, left half, or around 8,Fig. 4D, left half), suggesting that ABA did not stimulate the BSCs’ VHA in the dark.

However, when the bath medium pH (pH_EXT_) was elevated from its value of 5.6 (used so far in all our assays of pH_CYT_ in the light) to 7.5, more in keeping with the alkalinization of xylem sap observed in the dark (Grunwald et al., 2022), the BSCs reproduced the ABA-induced cytosol alkalinization (pH_CYT_ increased roughly from 7.7 to about 8.6; Fig. 4C, right, or to about 8.7, Fig. 4E, left). This pH_CYT_ response to ABA in the dark was abolished by bafilomycin A1 (Fig. 4E, right), confirming that the cytosol alkalization was due to VHA stimulation by ABA. Bafilomycin A1, by itself, did not affect the pH_CYT_ when pH_EXT_ was 7.5 (pH_CYT_ remained around 7.7; Fig. 4E), but in contrast, it acidified the cytosol when pH_EXT_ was 5.6 (pH_CYT_ declined from about 8.2 to 7.4; Fig. 4D), suggesting that in acidic medium, though not responding to ABA, VHA retained an inhibitable basal activity even in the dark.

The ABA-induced cytosol alkalinization (i.e., VHA stimulation) in the alkaline medium in the dark (Figs. 4F-4H) was abolished by adding BAPTA-AM (Fig. 4F), BAPTA-K_4_ (Fig. 4G) or La^3+^ (Fig. 4H) to the medium. This suggests that the ABA response (pH_CYT_ elevation), enabled by the external alkalinity, depended additionally on Ca^2+^ influx, very likely via Ca^2+^-permeable channels.

### The SKOR K^+-^efflux channel affects plant water management under drought

#### ABA-effect-mimicking ionic milieu composition enhances the BSCs’ K^+^-release channels

To test the hypothesis that the pH and [Ca^2+^]_CYT_ changes induced by ABA in and outside the BSCs enhance the transmembrane K^+^ transport, we examined the *in-situ* K^+^ channel activity in isolated BSCs protoplasts under eight combinations of ionic conditions corresponding to “resting” conditions, and to different levels of “ABA-effect”-like conditions, based on our experimental results: pH_EXT_ 5.6 and 7, pH_CYT_ 7.5 and 8, and free [Ca^2+^]_CYT_ of *nominally* about 100 nM and 600 nM, the two latter values similar to the ranges reported for GCs). (e.g. Irving et al., 1992; Allen et al., 1999).

We used patch clamp in the whole cell configuration and based our analyses on the records acquired after about 10 min elapsed since breaking into the cells, when the cell records seemed to have attained steady-state. The cell stability was aided by holding the cell membrane at -23 mV between the depolarizing or hyperpolarizing pulses. This holding potential (E_H_) drove a minimum of membrane current, usually close to 0.

We focused on the K^+^-release (K_OUT_) channels selecting them by their depolarization-dependent activity. By simple eyeballing and counting the cells with various current types, about 80 % of the BSCs from WT (Ler) plants were found to harbor a variable activity of the K_OUT_ channels, confirmed as such by their voltage-dependent and time-varying membrane currents and their reversal potentials (E_REV_). The mean E_REV_ of the K_OUT_ channels was -38 ± 10 mV (±SD, n=77), reflecting a prominent contribution of the only negative Nernst potential (E_ion_) of K^+^ (the E_ion_ values calculated for the potentially membrane-permeant ions in our experimental solutions were: E_K+_ -81 mV, E_Mg2+_ 0 mV, E_Cl-_ +21 mV, E_Ca2+_ +117 mV and E_H+_ between +29 mV (at pH_EXT_/pH_CYT_ 7/7.5) and +140 mV (at pH_EXT_/pH_CYT_ 5.6/8).

The activity of the K_OUT_ channels could be quantified in some cells by fitting the Boltzmann equation to the experimentally derived specific membrane conductance (G’) vs membrane potential (E_M_) relationship (as in Wigoda et al., 2017). To include also cases not amenable to Boltzman fitting, we quantified the K^+^ channel activity in all cases using the maximum specific membrane conductance for K^+^ (G’) attainable at the extreme edge of the used K_OUT_-activating E_M_ range (E_M_ of +97 to +117 mV) under the abovementioned eight different ionic millieu combinations. G’ was the highest at an ionic combination of pH_CYT_ 8, pH_EXT_ 7 and [Ca^2+^]_CYT_ =∼600 nM and the 2^nd^ highest at pH_CYT_ 7.5, pH_EXT_ 7 and [Ca^2+^]_CYT_

=∼600 nM (Fig. 5A).

#### The SKOR K^+-^release channel and plant water balance under drought

We tested whether the expected K^+^-deficiency of the *skor* plants leaves (Gaymard et al., 1998); Fig. 6C) affects their K_leaf_ compared to those of the WT plants. Indeed, while the K_leaf_ of detached leaves was on average similar in WT (Col) and in *skor* plants (about 30 mmol m^-2^ s^-1^ MPa^-1^; Fig. 5B, Suppl. S6), the leaves of the ABA-treated plants differed between the genotypes: ABA decreased the K_leaf_ in both WT and *skor* (similar to Shatil-Cohen et al., 2011), the K_leaf_ of the ABA-treated *skor* leaves declined less than K_leaf_ of the WT leaves (down to about 10 vs. 3 mmol m^-2^ s^-1^ MPa^-1^, respectively; Fig. 5B, Suppl. S6).

**Figure 6.**
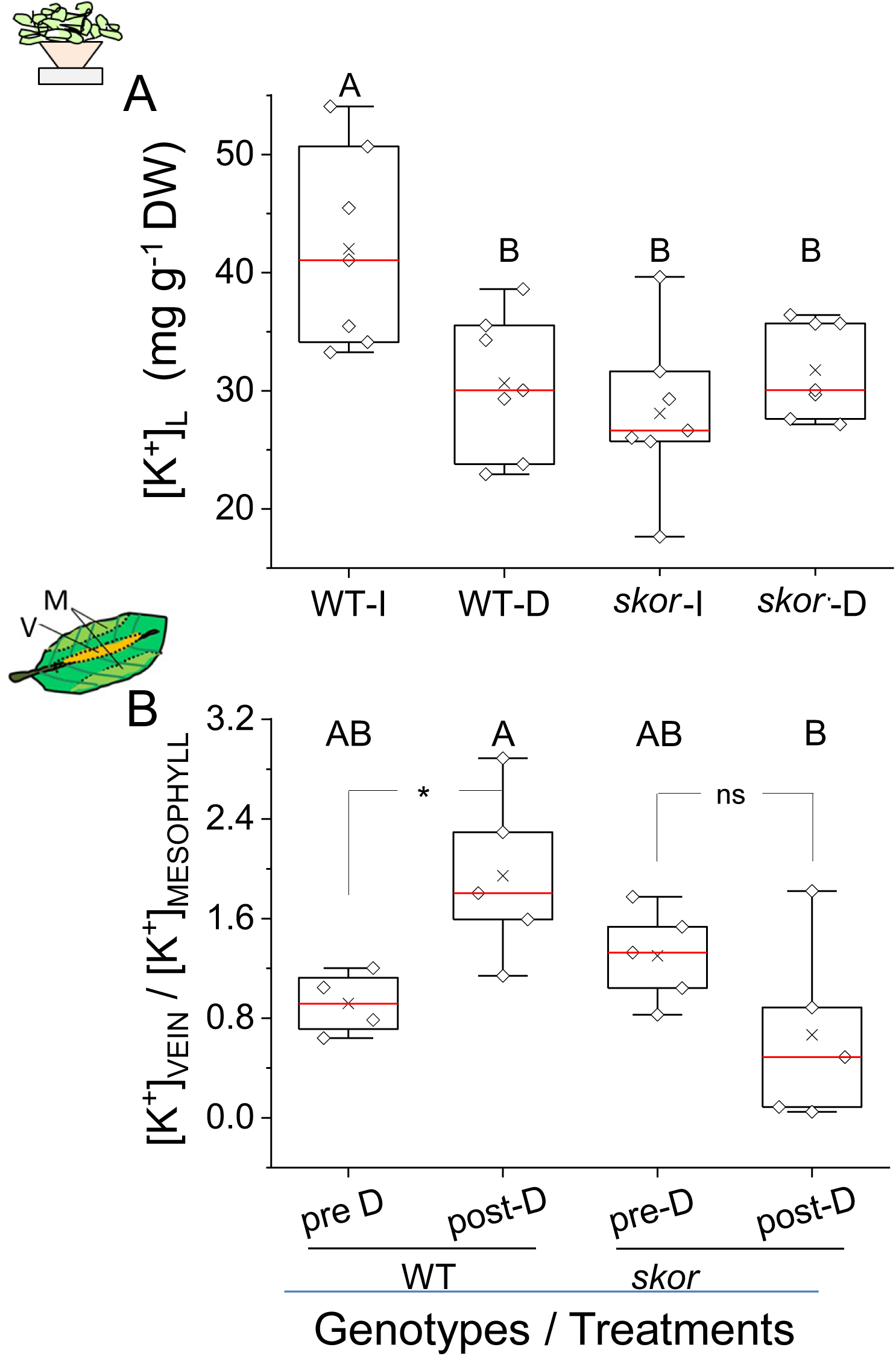
The concentration of K^+^ accumulated in the leaves ([K^+^]_L_) of WT and *skor* mutant plants exposed to water deprivation (D). **A.** [K^+^]_L_ determined by flame photometry in representative central rosette leaves at the end of the experiment of Fig. 5C. WT-I and *skor-I*: well-irrigated plants, WT-D, *skor-*D: droughted plants. Note the relatively large decline of [K^+^]_L_ due to drought in WT plants vs. the insignificant change of [K^+^]_L_ in the *skor* mutant plants. **Inset.** A schematics of leaves in a four-plant pot in the greenhouse. **B.** The ratio of [K^+^] determined by ICP-EOS in different leaf parts sampled on the same plants before and after 2 weeks of water deprivation in an experiment similar to A: “Vein” – a mid-rib-vein-containing leaf part, presumably representing mainly the vascular tissue, and “Mesophyll” – a leaf edge with presumably mainly mesophyll tissue. *: a significant difference (P=0.02423, by non-paired t-test on the ratios, *two*-tailed, equal variance), or P= 0.01242 (by a similar t-test on *log10* of the ratios), ns: not significant. Note the drought-induced *increase* of vein / mesophyll ratio in WT leaf fragments, and the absence of such an increase in *skor*. **Inset.** A schematics of the sampled leaf parts. See also **Suppl. Fig. S8**.

These results led us to *predict* that water loss of whole *skor* plants stressed by water deprivation will be greater than that of the WT plants. To test this, we grew both WT (Col) and *skor* (Col) plants in pots in a greenhouse (Fig. 5C) where half of each group was subjected to drought treatment for 13 days, followed by 8 days of re-irrigation. After 8 days of water deprivation, when the soil water content decreased below 30 % of full capacity, the daily as well as momentary normalized water loss by transpiration in *skor* mutant plants was higher than in WT plants (Fig. 5F, Suppl. Fig. S7B, respectively), despite a similar soil water content in all pots throughout the drought period (Fig. 5E). Even during the eight days of restored irrigation, the normalized daily transpiration of skor plants remained higher than in the WT. In contrast, under uninterrupted water sufficiency, the two genotypes showed no difference in normalized daily transpiration during this period (Fig. 5F), as if the activation of SKOR by ABA (in the droughted WT) contributed to water conservation.

Since the prominent effect of SKOR mutation was the diminished K^+^ accumuation in the shoot (Gaymard et al., 1998), we set out to explore a possible link between the *skor*’s altered water management under water defficiency stress with its K^+^ management in these conditions. As expected, among the well-irrigated plants, at the end of the experiment, [K^+^] in the central rosette leaves ([K^+^]_L_) of *skor* plants was lower than [K^+^]_L_ in WT plants (only 67 % of WT; Fig. 6A). At the same time, among plants which underwent a two-weeks drought (and 8 days of re-irrigation), the [K^+^]_L_ in the WT plants was appreciably lower than WT in the former group (down to 71 % of the well-irrigated WT; Fig. 6A left). However, [K^+^]_L_ in the *skor* plants remained unaltered (Fig. 6A right). The loss of K^+^ from the droughted WT leaves *is consitent* with the abovementioned ABA-induced K^+^ release from the BSCs via K_OUT_ channels into the apoplast and K^+^ removal from the leaf through the BSCs then via xylem and/or phloem. The lack of such a decline in droughted skor leaves *is consistent* with SKOR being the K_OUT_ channel involved. In plants treated similarly in an independent experiment, [K^+^] differed between different parts of leaves sampled from the same plants before and after drought (Suppl. Fig. S8). [K^+^]_V_, [K^+^] in the midrib vein, assumed to represent mainly the vasculature (including the bundle sheath) was compared to [K^+^]_M_, [K^+^] in the leaf edges, assumed to represent mainly mesophyll (Fig. 6B, inset). The ratio of [K^+^] ratio of [K^+^]_V_ to [K^+^]_M_ in the WT leaf increased following water deprivation while in the *skor* leaf no increase was observed after a parallel treatment (Fig. 6B). The increased [K^+^]_V_ / [K^+^]_M_ ratio in the WT leaf vs.its invariability in the *skor* leaf *is –* again *– consistent* with ABA-induced K^+^ release through and from the BSCs, via SKOR channels, into the leaf vasculature.

### The leaf anatomy, [K^+^]_L_ and K_leaf_

The similarity between the well-irrigated WT and *skor* plants in term of of K_leaf_ and plant normalized transpiration (daily, Fig. 5F or momentary, Suppl. Fig. S7B) could not be predicted based on the comparison of their leaves anatomy: the vein density of the skor leaf is higher than that of the WT (by 16 %; Suppl. Fig. S9A). However, while stomata index on the adaxial (top) leaf side is similar in both *skor* and the WT, the skor’s abaxial stomata index is lower than in WT (by 11 %; Suppl. Fig. S9B) and the stomata density of *skor* is also lower than that of the WT (on the abaxial side: lower by 16 % and on the adaxial side – lower by 14%).

## DISCUSSION

Our results can be summarized in terms of a scheme (Fig. 7) of two counteracting signal transduction pathways within the BSCs, (a) the activation by BL of the AHA2 proton pump and the resulting increase in plant transpiration, and (b): the dominant inhibition of AHA2 by ABA and the resulting decline in plant transpiration.

**FIGURE 7.**
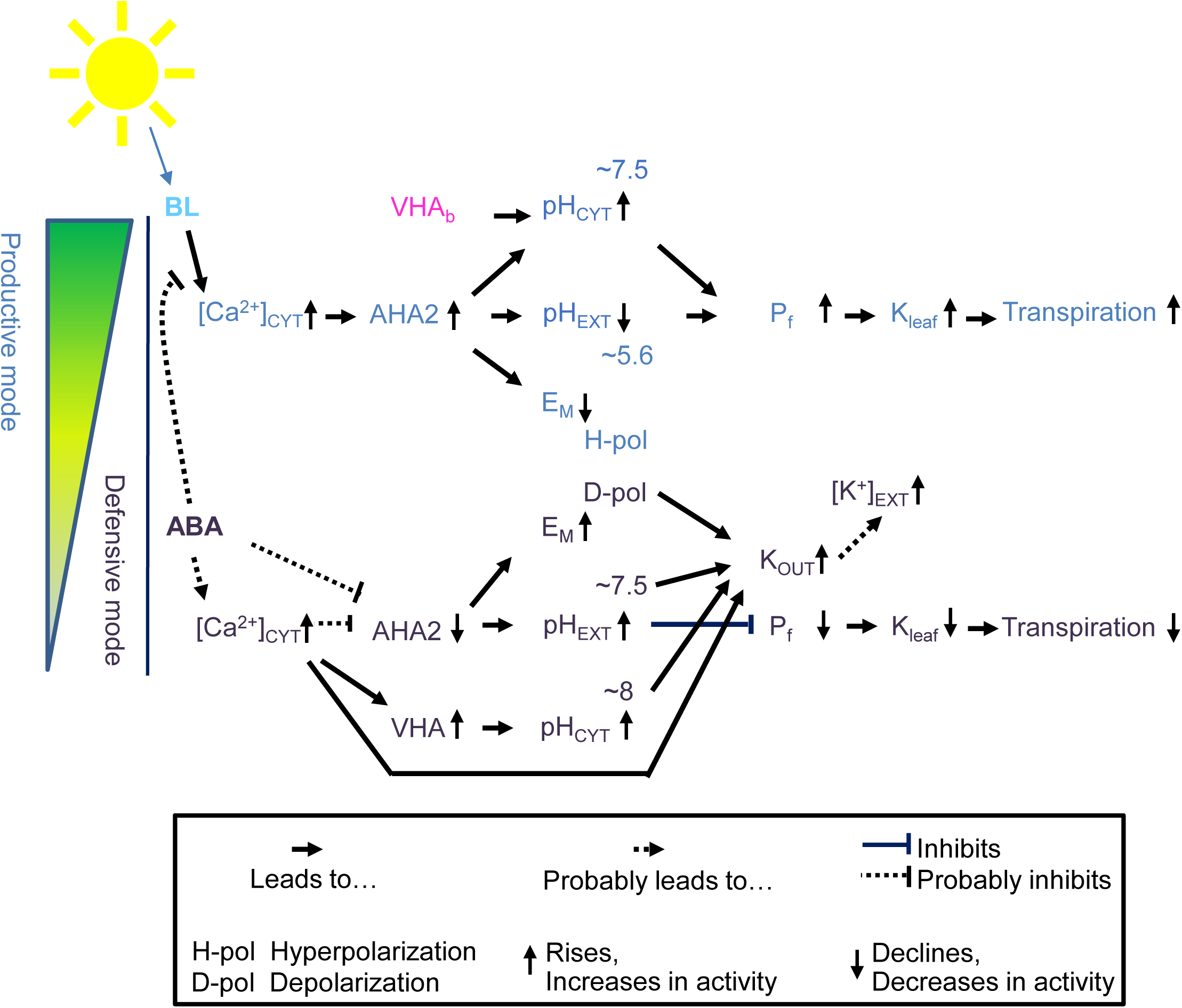
A schematic portrayal of the two signaling cascades within the BSCs. Blue light (BL) and the abscisic acid (ABA) exert their effects on K_leaf_ and – eventually – on transpiration, via H^+^-pump regulation, pH changes and modulation of AQP activity and possibly also of the activity of K_OUT_ channels.

### Blue light (BL) activation of the BSCs’ AHA2 and K_leaf_ is mediated by Ca^2+^

Recently, we established linkages between the BL-induced phot-mediated AHA2 activation within the BSCs, the BSCs hyperpolarization, the acidification of the xylem perfusion solution (XPS) in a detached leaf (pH_EXT_ decrease), and the resulting increase of the leaf hydraulic conductance, K_leaf_ (Grunwald et al., 2021, 2022). Here we demonstrate that this BL-induced activation of the BSCs’ AHA2 requires elevated [Ca^2+^]_CYT_. This conclusion stems (a) from decreasing K_leaf_ in detached leaves by the cytosolic Ca^2+^ chelator BAPTA-AM (Fig. 1A), (b) the inhibition of AHA2 activity in BSC protoplasts by prevention of the BL-induced [Ca^2+^]_CYT_ increase, either directly by BAPTA-AM (Fig. 1D), or by decreasing the [Ca^2+^]-gradient-driven influx of Ca^2+^ from outside by BAPTA-K_4_ (Fig. 1B), which likely occurs via Ca^2+^-permeable La^3+^-blockable channels (Fig. 1C). The scenario of phot-mediated activation of Ca^2+^ channels in the plasma membrane by BL and a consequent activation of AHA2 – manifested as a BSC hyperpolarization – is suggested schematically also in our model, by blue arrows 1-3 (Figs. 11A, 11B).

BL-induced activation of AHA via elevated [Ca^2+^]_CYT_ was also reported in other systems: (a) Ca^2+^ channels-mediated BL-induced Ca^2+^ influx and an increase of [Ca^2+^]_CYT_ in Arabidopsis seedlings (Babourina et al., 2002) and (b) phot receptors-mediated BL activation of Ca^2+^ channels in Arabidopsis mesophyll cells (Stoelzle et al., 2003; Harada et al., 2003)). In the BSCs, among the plasma membrane-residing Ca^2+^ channel candidates could be, for example, some of the Ca^2+^-permeable CNGCs (cyclic-nucleotide-gated channels; reviewed by Dietrich et al., 2020), 17 of which figure in the BSCs transcriptome (Wigoda et al., 2017), or the GLR channels (Glutamate Receptor channels; reviewed by Grenzi et al., 2022); 18 of which are also expressed in the BSCs; ibid.). However, the molecular identity of the specific BL-activated BSCs’ Ca^2+^ channels remains to be established (Kleist and Wudick, 2022). Also, notably, our results do not eliminate an additional potential contribution of a “Ca^2+^-induced Ca^2+^ release” from internal Ca^2+^ stores, such as occurs from the vacuole during stomata closure via the AtTPC1 (a.k.a. SV channel; (Ward and Schroeder, 1994; Ye et al., 2021), a two-pore channel 1, the product of At4g03560), that is also expressed in the BSCs (Wigoda et al., 2017).

A link between the BL-induced [Ca^2+^]_CYT_ increase and AHA2 activation could be provided by a Ca^2+^-sensor-kinase pair, AtCBL7 (Calcineurin B-like 7, a.k.a. SCaBP3) and AtCIPK11 (CBL-interacting protein kinase11, a.k.a. PKS5), such as has been described in Arabidopsis seedlings roots challenged with a saline-alkaline stress (Yang et al., 2019) and both of which are expressed also in the BSCs; Wigoda et al., 2017. We thus speculate (modeling after Yang et al., 2019), that in BSCs the BL-induced [Ca^2+^]_CYT_ elevation dis-inhibits and activates the AHA2 by relieving its interaction with a CBL7/CIPK11 pair. Future assays will resolve whether such a scenario underlies the Ca^2+^-dependent BL-induced activation of the BSCs’ AHA2, which acidifes the BSCs’ apoplast, i.e., the xylem sap, which, in turn, activates AQPs and increases K_leaf_ (Grunwald et al., 2021; Grunwald et al., 2022; Fig. 1).

### The BSCs’ AHA2 is the target of the ABA inhibitory signaling

Under stress, ABA concentration raises and overcomes the BL signal FIGtransduction, transferring the plant from a productive to a defensive mode (Fig. 7). Based on our earlier demonstration of the AHA2’s prominent role in the leaf xylem sap acidification (Grunwald et al., 2021, 2022), we now identify AHA2 as the BSCs-specific target of the ABA-inducible signaling cascade. This resembles the Arabidopsis seedlings hypocotyl (Hayashi et al., 2014), in which ABA inhibited AHA2 (Hayashi et al., 2014) and differs from the Arabidopsis guard cells, in which ABA inhibited AHA1 (Hayashi et al., 2011). Our conclusion stems from (a) the absence of the ABA response (xylem perfusate alkalinization) in the detached leaves of the *aha2-4* mutant plants (Fig. 2B) practically devoid of AHA2 activity (Haruta, M., Burch, H.L., Nelson, R.B., Barrett-Wilt, G., Kline, K.G., Mohsin, S.B., Young, J.C., Otegui, M.S., and Sussman, 2010; Haruta and Sussman, 2012), (b) the restoration of the ABA response in the leaf of *aha2-4* complemented with the BSCs-directed *SCR:AHA2* (Grunwald et al., 2021; Fig. 2C) and (c) the absence of the ABA-induced depolarization in single BSCs from the *aha2-4* mutant plants and its restoration in BSCs from the *SCR:AHA2-*complemented *aha2-4* plants (Figs. 2D-2F). The disruption of the above ABA-induced responses by the mutant abi-1 (Figs. 2G and Suppl. Fig. S3) suggests a similarity of the BSCs’ and the GCs’ ABA early signaling sequences; in both systems, the target AHA inhibition requires an intact ABI1, capable of being sequestred by the ABA-bound receptor complex and pulled out of circulation (Leung, J., Merlot, S., and Giraudat, 1997; Merlot et al., 2001; Imes, D., Mumm, P., Böhm, J., Al-Rasheid, K. A.S., Marten, I., Geiger, D., and Hedrich, 2013) . Thus in both systems, enabling the downstream phosphorylation-dependent reactions is necessary for AHA inhibition.

### Do ABA-activated Ca^2+^ and/ or anion channels contribute to the ABA-induced BSCs depolarization?

Early depolarization, due to the channel-mediated influx of Ca^2+^ ions and/ or efflux of anions, are both hallmarks of an early ABA effect on guard cells (Schroeder and Hagiwara, 1990; Roelfsema et al., 2004). In Arabidopsis suspension cells, the ABA-induced stable depolarization – also attained within a few minutes – included both activation of anion efflux and proton pumping inhibition, which were both Ca^2+^ dependent (Brault et al., 2004).

In the BSCs, to our surprise, we were unable to observe an early depolarization during the first 5 min after the exposure to ABA (Suppl. Figs. S2A,C) although we did observe a depolarization during minutes 10-15 (Suppl. Figs. S2B,D) and demonstrated that this “late” BSCs depolarization is due to AHA2 inhibition (Figs. 2A-F). This apparent lack of channel-mediated depolarization is not likly to be due to the lack of the relevant channels; in addition to the abovementioned BSCs candidate Ca^2+^ channels, also 3 out of the 4 Arabidopsis homologues of the slow-activating anion channel SLAC1 (SLAH1 to SLAH3) were found expressed in the BSCs (Wigoda et al., 2017), even if the major contributor to guard cell depolarization, SLAC1 itself, was missing (or at least, was not detected) in the BSCs transcriptome (ibid.). SLAH3, although proclaimed to be 20 times less permeable to Cl^-^ than to NO^3-^ (Geiger, D., Maierhofer, T., Al-Rasheid, K. A S., Scherzer, S., Mumm, P., Liese, A., Ache, P., Wellmann, C., Marten, I., Grill, E., Romeis, T., and Hedrich, 2011), subserved Cl^-^ efflux from Arabidopsis root epidermal cells which caused a small fast depolarization within five minutes of an ABA treatment (Planes et al., 2015). How then could such an early depolarization of the BSCs (due to both Ca^2+^ influx and anion efflux) be absent in the BSCs? Could it be overcome by a co-evoked efflux of K^+^ via ABA-activated BSCs’ K_OUT_ channels?

We suggest that the primary cause of the delayed depolarization appearing only at min 10-15 after the exposure of the BSC protoplast to ABA (Suppl. Fig. S2B), which we show to result from AHA2 inhibition (Figs. 2D-F), is the imbalance of the positive charges resulting from the *cessation* of proton extrusion by the AHA2. The carriers of the – now unbalanced – net positive charges are very likely the BSCs’ plasmalemmal H^+^ symporters, such as an H^+^/NO_3_^-^ symporter and an H^+^/ K^+^ symporter. This notion is supported by our earlier experiments, when we enhanced the XPS alkalinization (i.e., escape of protons from the xylem to the adjacent BSCs) by adding 10 mM KNO_3_ to the XPS (Grunwald et al., 2021). In the BSCs transcriptome these carriers are represented, respectively, by AtNRT1.1 (Fang et al., 2016; Wigoda et al., 2017) and by the K^+^ uptake permeases, AtKT2 and AtKUP11 (Wigoda et al., 2017) which are likely to be H^+^-coupled (Elumalai et al., 2002; Ahn et al., 2004).

### Potential mediators of AHA2 inhibition by ABA

#### ABA-induced cytosol alkalinization is not a signal for AHA2 inhibition

ABA-elevation of [Ca^2+^]_CYT_ and stomata closure were both conditional on a rapid cytosol alkalinization in *Nicotiana tabacum* GCs (Li et al., 2021). In contrast, in Arabidopsis roots, ABA did not trigger rapid cytosolic Ca^2+^ or H^+^ dynamics (Waadt et al., 2020). Since in the BSCs ABA activated the VHA and alkalinized the cytosol within the first few minutes of its appplication (Suppl. Fig., S4 and see more below), could the resulting cytosolic alkalinization be the primary reason for the cessation of AHA2 activity (as, for example, in Hoffmann et al., 2019)? We reject this notion, since while the cytosol-alkalinizing VHA activity was abolished by bafilomycin A1 and ABA lowered the cytosol pH (Fig. 3C, right), the ABA-induced AHA2 inhibition, manifested as ABA-induced BSCs depolarization, remained unaffected (Suppl. Figs. S2D and S2B). A further support may be derived from our guiding model system, the GCs, wherein cytosol alkalinization was required but not sufficient for stomata closure (Irving et al., 1992).

#### Elevation of the free cytosolic Ca^2+^ concentration

ABA-induced Ca^2+^ signals are common in guard cells (e.g., Kinoshita et al., 1995), even if they appear only in about 75% of Arabidopsis GCs (Huang et al., 2019), and some of the guard cell models show Ca^2+^ as mediating the ABA-induced inhibition of the GC’s AHA (e.g., (e.g., (Chaves et al., 2011; Cai et al., 2017; Albert et al., 2017)).

In the BSCs, increased [Ca^2+^]_CYT_ mediated the *activation* of AHA2 by BL (Figs. 1B-1D), while the ABA-induced *inhibition* of the BSCs’ AHA2 overcame this BL-induced activation (Figs. 2A,2D, Suppl. Figs. S2B, S3A, S3C). This ‘overruling of BL by ABA’ occurred by an as yet unresolved mechanism, one of which *could* be ABA-induced elevation of [Ca^2+^]_CYT_ (see our scheme, Fig. 7) and the detailed BSC model, Figs. 8C, 8D).

**Figure 8.**
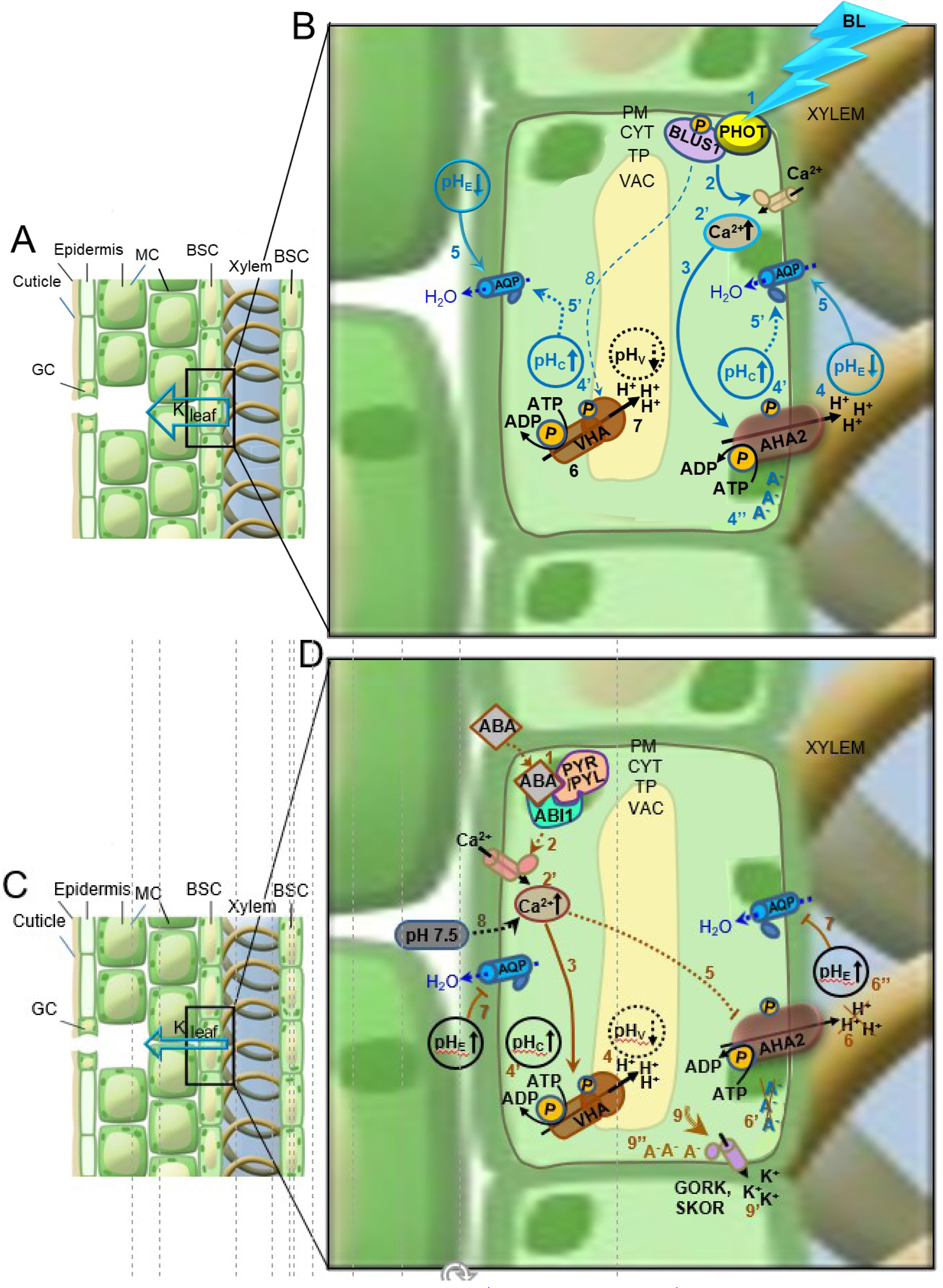
Proposed BSCs-autonomous BL and ABA signal transduction sequences. An artist’s rendering of changes in an ionic balance between the BSCs and the xylem, due to a simultaneous regulation of two H^+^-ATPases – the plasma membrane (PM) AHA2 and the tonoplast (TP) VHA, the K^+^-efflux channels, GORK and SKOR, and the PM aquaporins. **A, C.** A leaf cross-section. GC, a stomata guard cell; MC, mesophyll cell; BSC, bundle sheath cell; K_leaf_, the leaf hydraulic conductance. Note the different sizes of the K_leaf_ arrow, indicating different K_leaf_ levels. **B, D.** A BSC in an expanded view. PM: plasma membrane, CYT: cytosol, TP: tonoplast, VAC: vacuole. **B. The Blue-Light (BL) signaling pathway**; BL perception (**1**) by the phototropin receptors complex (PHOT+BLUS1) leads to the activation of PM Ca^2+^ channels (➔, **2**)^a^, enabling influx of Ca^2+^ from outside the cell and increasing [Ca^2+^]_CYT_ (**2’**), thereby **activating AHA2** (➔, **3**)^b^, ultimately resulting in H^+^ extrusion and xylem sap acidification (**4**)^b^, cytosol alkalinization (**4’**)^b^ and in hyperpolarization (**4’’**)^b, c^. Acidic xylem sap activates aquaporins (➔**, 5**)^d^, presumably, when combined with alkaline cytosol already known to activate AQPs (**^….^>, 5’**)^e^. **VHA is basally active at pH_EXT_ 5.6** while in the DARK (**6**)^f^, contributing to the pH_CYT_ alkalinity (➔**, 4’**), presumably by extruding H^+^ into the vacuole (➔**, 7**)^g^. VHA was *reportedly* activated by BL (^….^>, 8)^ii^, but in our work VHA basal activity was unaffected by adding BL to RL (➔, 7)^h^, nor by the lowering of the [Ca^2+^]_CYT_ level^i^. Alkaline cytosol activates AQPs (**^….^>, 5’**)^e^, however, presumably only when combined with acidic xylem sap (➔**, 5**)^d^ and this combination of pH_EXT_ and pH_CYT_ (**4, 4’**) underlies the BL-increased K_leaf_^d^, which also depends on increased [Ca^2+^]_CY_ ^cc^. **D. The ABA signaling pathway.** ABA binds to intracellular receptors^k^ recruiting the PP2C phosphatase ABI1 to the complex (**1**)^l^. ABI1 mediates the ABA-induced AHA2 inhibition^m^ as follows: ABA (via the receptor complex) activates Ca^2+^ channels in the plasma membrane (^….^**>**, **2**)^n^, thus elevating [Ca^2+^]_CYT_ (Ca^2**+**^↑, **2’**)°. ABA stimulates the VHA (^….^**>, 3**) which extrudes protons across the TP acidifying the vacuole (pH_v_↓, **4**)^g^ and, concomitantly, alkalinizing the cytosol (pH_c_↑, **4’**)^p^, which is VHA specific^q^ and [Ca^2+^]_CYT_ dependent**^r^**. On the background of RL+BL, at pH_EXT_ 5.6, [Ca^2+^]_CYT_ elevated by ABA° and /or pH_CYT_ elevated by ABA-stimulated VHA ^p^, inhibit the AHA2 (^….^I, **5)^s^,** (^….^I, **5’**)**^ss^**, which halts proton extrusion via the PM (**6**), consequently depolarizing the BSCs (**6’**)^t^ and alkalinizing the xylem sap (pH_e_ ↑, **7**)^u^. ABA-induced xylem alkalinization inhibits aquaporin activity (**7**)**^d^**. In the DARK, at pH_EXT_ 5.6, VHA does not respond to ABA^v^, but at pH_EXT_ 7.5 ABA does elevate pH_CYT_ (pH_c_ ↑, **3**)**^rr^** depending on Ca^2+^ influx (^….^>**8**)^r^. Furthermore, ABA-generated conditions activate the GORK/SKOR K^+^-efflux channels (**9**)^w^, enabling K^+^ efflux, and, potentially, increasing the [K^+^] in the xylem (**9’**)**^x^**.

How could *both* AHA2 activation and its inhibition operate via a rise in [Ca^2+^]_CYT_? We speculate that these contrasting responses originate in two signaling pathways which generate different [Ca^2+^]_CYT_ signatures in the BSCs, such as shown multiple times elsewhere (e.g., McAinsh et al., 1995; a review by (McAinsh and Pittman, 2009). Thus, a specific Ca^2+^ signature could mediate the BL-induced AHA2 *activation*, already described above by evoking the interaction of the BSCs’ AtCBL7 with CIPK11 (modeled after Yang et al., 2019). We speculate further that a different, ABA-generated, [Ca^2+^]_CYT_ signature underlies the BSCs’ AHA2 *inhibition*, mediated by an interaction between the Ca^2+^ sensor AtCBL2 (which the BSCs also express; Wigoda et al., 2017) and its dependent kinase, CIPK11 ( “the servant of two masters” in both scenarios). We model this sequence after an alkaline-stress-induced AHA2 inhibition (in Arabidopsis seedling roots), wherein the AtCBL2-CIPK11 pair transduced a [Ca^2+^]_CYT_ elevation into a phosphorylation of the AHA2-inhibitory site, SER^931^ (Fuglsang et al., 2007).

For completeness, our scheme (Fig. 7) includes an alternative hypothetical pathway, modeled on GCs, of ABA-induced disruption of the BL signaling: by ABA-generated phosphatidic acid (Takemiya et al., 2006) and / or by ABA-activated SnRK2 kinase, OST1 (Open STomata1; Hayashi et al., 2011). A definitive resolution of this pathway validity in the BSCs awaits further exepriments.

### In the BSCs, VHA is a target of ABA-induced stimulation

Unlike in the Arabidopsis GCs, where ABA has been recently reported to activate transiently the AHA2 thereby transiently alkalinizing the cytosol (Pei et al., 2022), the ABA-induced BSCs cytosol alkalinization that we observed in the BScs, could not be due to any AHA, including AHA2 (Fig. 3A) and the tonoplast-localized P-type group III H^+^-ATPase AHA10 (At1g17260; Appelhagen et al., 2015; Kriegel et al., 2015; also expressed in the BSCs; Wigoda et al., 2017)), all of which we eliminated using vanadate (Fig. 3B). The remaining candidates for the ABA-induced cytosol alkalinization in the presence of vanadate were the multi-subunit V-type H^+^-ATPase, VHA (Schumacher and Krebs, 2010) and the homo-dimeric V-PPase, AVP1 (AT1G15690; Segami et al., 2018). Both the VHA and the AVP1 (VHP1) were implicated in the vacuole acidification and concomitant cytosol alkalinization in the guard cells of Arabidopsis and *Vicia faba* and both were required together for a normal ABA-induced stomata closure in mature Arabidopsis rosettes (Bak et al., 2013)). However, the constitutive expression of AVP1 in the double VHA mutant *vha-a2 vha-a3* could *not* compensate for VHA deficits in tissues *other than guard cells* (Segami et al., 2018; Kriegel et al, 2015). Moreover, also our own demonstration in single isolated BSCs that a selective elimination of the BSCs’ VHA pump activity – either by its specific inhibitor bafilomycin A1 (Muroi et al., 1994) Fig. 3C) or by the double *vha-a2 vha-a3* mutation (Fig. 3D) – abolished completely the ABA-induced pH_CYT_ elevation, suggests that, in contrast to GCs, in the BSCs, VHA alone – and not their AVP1 (nor AHA10) – appears to be activated as a target of the ABA fast-signaling pathway, alkalinizing the cytosol within 0-5 min (Suppl. Fig. S4D). Notwithstanding, we have not ruled out that an ABA-insensitive background AVP1 activity is responsible, at least partially, for the elevated pH_CYT_ of the BSCs of the *vha-a2 vha-a3* double-mutant (Fig. 3D).

### The regulation of VHA

VHA is known to be regulated on a rapid time scale (minutes, e.g., Dechant et al., 2010) by dis- and re-assembly of its many subunits (Kane, 2012) , some of which, at least, is regulated by phosphorylation (reviewed by Wang et al., 2021; Seidel, 2022), although this has not yet been proven in plants (Seidel, 2022).

#### Does the BSCs’ VHA basal activity require light? VHA response to ABA – does

The inhibition of the AHAs by vanadate alone in the light (RL+BL), should have acidified the cytosol by the now-unbalanced proton leak into the cytosol via the H^+^/K^+^ or H^+^/anion symport from the standard bath solution at pH_EXT_ 5.6 and perhaps also from the acidic vacuole. The unexpected *lack* of pH_CYT_ decline, and even a small alkalinization (Fig. 3B, -ABA/+VAN, compared to -ABA/-VAN), suggests a compensatory basal VHA activity overcoming the proton leak. Consistent with this notion, pH_CYT_ declined when both pumps were inactivated: AHA2 by darkness (Grunwald et al., 2022) and VHA by bafilomycin A1 (applied by itself; Fig.4D, -ABA/+BAF vs. -ABA/-BAF). Further, suggesting that VHA is basally active both in the dark and in the light, is the lack of change of pH_CYT_ under the transition from RL to RL+BL (Fig. 4A, left), in stark contrast to the activation of AHA2 by RL+BL, compared to dark or to RL (Grunwald et al., 2022, their Fig. 5). Alternatively, rather than indifferent to light, the basal BSCs’ VHA activity could be *overly sensitive* to BL (more so than AHA2), such that the very brief episodic BL illumination used for seeking the GFP-labeled BSCs, could have sufficed to maintain the VHA at a basal activity level throughout the dark or RL treatments. VHA activation by dark-to light transition has been reported for other plant cell types (Dietz et al., 1998; Klychnikov, O.I., Li, K.W., Lill, H., and De Boer, 2007).

Regardless of VHA sensitivity to BL, the *stimulation* of VHA by ABA *did* require light, manifested as ABA-induced cytosol alkalinization under RL+BL (Fig.4B, left), on the one hand, and, on the other hand, as the lack of the ABA response in the dark, in the same standard bath solution at pH_EXT_ 5.6 (Figs. 4C- left, 4D-left).

#### VHA regulation by [Ca^2+^]CYT

Interestingly, on the backgroud of vanadate-inhibited AHAs, the BSCs’ VHA *basal* activity in the light is not sensitive to the *decline* of [Ca^2+^]_CYT_, as suggested by the lack of the effect of BAPTA-AM, added by itself, on the BSCs’ pH_CYT_ (Fig. 4B). In contrast, VHA *stimulation* by ABA in the light (RL+BL) does require at least a “resting” level of [Ca^2+^]_CYT_ (or such as BL confers), as can be concluded from the BAPTA-AM abolishment of the ABA-induced cytosol alkalinization (Fig. 4B). Moreover, the response of VHA to ABA in the dark, which was restored in an alkaline medium (Figs. 4C, right and 4E-4H, left), requires an elevated [Ca^2+^]_CYT_, as can be deduced from the abolishment of the ABA response upon the decrease of Ca^2+^ influx from the bath (Figs. 4G, 4H), which replicated the direct chelation of the cytosolic Ca^2+^ (Fig. 4F).

It remains yet to be resolved which of the two signals that together stimulate VHA at pH_EXT_ 5.6 – the light (RL+BL) or ABA, or both – elicits the increase of [Ca^2+^]_CYT_ which mediates the cytosol alkalinization. Additionally, it remains yet to be resolved which of the two signals that together stimulate VHA in the dark – the alkaline medium or ABA, or both – elevates the [Ca^2+^]_CYT_ – a prerequisit for cytosol alkalinization.

Notwithstanding the exact identitity of the inducing signal, we now speculate that stimulation of the BSCs’ VHA activity by [Ca^2+^]_CYT_ elevation is mediated by Ca^2+^ sensors, such as the tonoplast-associated CBL2,CBL3 shown to increase VHA activity (Tang et al., 2012) through their associated (as yet unspecified) CIPKs (reviewed by Saito and Uozumi, 2020). Both CBLs are expressed in the BSCs (Wigoda et al., 2017).

### From the interplay between the BSCs’ pumps to the regulation of plant transpiration under drought

The balanced regulation of the cytosolic pH based on the activity of proton pumps in the cytosol-delimiting membranes was reviewed recently (Sze and Chanroj, 2018; Cosse and Seidel, 2021). Both the BSCs’ pH_CYT_ and pH_EXT_ are strongly influenced by the interplay between the two BSCs’ H^+^-pumps and their regulation by the BL and ABA signaling pathways. We now propose that the pH balance within the two compartments – the cytosol and the xylem sap – is a major mechanism underlying the fast regulation of the activity of the BSCs’ plasma membrane aquaporins by BL and by ABA, and through this, the fast regulation of the whole plant transpiration. We rely on the following evidence:

a. Both, BL and ABA alkalinize the BSCs’ cytosol (as shown, respectively, in Grunwald et al., 2022 and repeatedly here). Cytosol alkalinization, in turn, promotes aquaporin activity via a histidine (H197 in Arabidopsis) in their cytosolic loop D, conserved in all PIP aquaporins (Tournaire-Roux et al., 2003).
b. In contrast, BL and ABA cause diametrically opposite changes in the the xylem sap: BL acidifies, while ABA alkalinizes it (as shown again, respectively, in Grunwald et al., 2022 and here).
c. We have recently demonstrated an inverse link between the BSCs osmotic water permeability, P_f_, and pH_EXT_: low pH_EXT_ increases P_f_ while high pH_EXT_ diminishes P_f_ (Grunwald et al., 2021). Through these pH changes, we linked BL with the increase of the BSCs P_f_ (Grunwald et al., 2022) and ABA – with the decrease of the BSCs P_f_ (Shatil-Cohen et al., 2011).
d. Further, we linked the changes of the BSCs P_f_ with the changes in the leaf hydraulic water conductivity, K_leaf_, both under BL and under ABA (Grunwald et al., 2022; Shatil-Cohen et al., 2011).
e. These links indicate that the high P_f_ level – under BL – reflects the *coincidence* of alkalinity in the cytosol (due mainly to the AHA2 activity, and maybe, also to some basal VHA activity) with external acidity (also due mainly to the AHA2 activity). In lieu of such coincidence – under ABA – when the pH_EXT_ is high (because of AHA2 inhibition), even if pH_CYT_ is also high (due to VHA stimulation), the P_f_ level is low (Shatil-Cohen et al., 2011, Grunwald et al., 2021) and thus K_leaf_ is low (Fig. 5B, left).

### The BSCs KOUT channels may play a role in the plant drought response

The finding that under ABA-effect-mimicking ionic conditions the K_OUT_ channels, which could potentially be activated in approx. 80 % of the BSCs, were most active under the combination of pH_EXT_ 7, pH_CYT_ 8 and free [Ca^2+^]_CYT_ around 600 nM (Fig. 5A) suggests that, like in the GCs, a drought signal mediated by ABA may cause a release of K^+^ into the apoplast (here – the xylem and/or further into the phloem). Of note, in the BSCs, K^+^ release could be mediated by any of the two K_OUT_ channels, SKOR and / or GORK expressed in the BSCs (Wigoda et al., 2017), separately or in heteromers (this is in contrast to GCs, which express only GORK). The slightly ameliorated ABA-induced decline of K_leaf_ of detached leaves of the *skor* mutant as compared to ABA-treated WT leaves (Fig. 5B, see also Suppl. Fig. S6), correlated well with the relatively higher normalized daily and momentary transpiration of water-deprived whole *skor* plants as compared to WT plants (Figs. 5F, Suppl. Fig. S7B, respectively). This suggests the involvement of SKOR in plant water conservation under water deprivation – perhaps by enabling K^+^ release from BSCs into the xylem.

### Physiological significance of K^+^ for leaf hydraulics

Recent report demonstrated the importance of K^+^ level in the plant to the plant water and carbon management in two dicot species, an Arabidopsis relative, rapeseed (*Brassica napus* L.) and cucumber (*Cucumis sativus* L.), and in two monocots, rice (*Oryza sativa* L.) and wheat (*Triticum aestivum* L.). In these plants, a decreased K^+^ accumulation in the shoot (attained by a long-term K^+^ starvation) decreased the leaf hydraulic conductance. In addition, the above dicots tended to show a decreased minor vein density (Lu et al., 2019). In spite of its lower than WT’s [K^+^]_L_ (Fig. 6, Suppl. Fig. S8), *skor* does not seem to conform to the label of ”K^+^-starved” plant, since, in contrast to those dicots, its vein density is higher than the WT’s (Suppl. Fig. S9). At any rate, the combination of anatomical traits renders the water dynamics of *skor* similar to WT’s when both are well irrigated and cannot explain the short-term difference in water dynamics under drought.

#### Is K^+^ release involved in plant response to drought stress?

Our results support a *causative* link between the observed decline of whole plant transpiration under drought and the observed decline of K_leaf_ of a leaf imbibed with ABA, on one hand, and the observed decline of the osmotic water permeability of the BSCs, P_f_, on the other hand. The intermediate elements of this link consist of the imposed or ABA-induced external alkalinity that we observed in a whole leaf and the still hypothetical ABA-induced K^+^ release via the BSCs K_OUT_ channels. It is worth noting, however, that such ABA-induced K^+^ release through and from the BSCs, via SKOR channels into the leaf vasculature may underlie the differences between WT and *skor* in their K^+^ management under drought (Fig. 6 and Suppl. Fig. S8). Moreover, ABA-induced loss of K^+^ from the BSCs may allow K^+^ transport towards the roots via the phloem and explain, in part, the observation that under water deficiency, growth is readily inhibited and the growth of roots is favoured over that of leaves (Hsiao and Xu, 2000). On the other hand, the increase of K^+^ concentration in the xylem would decrease the osmotic pressure of the xylem sap (for example, by 10 kPa upon an increase from 0.5 mM KCl to 2.5 mM KCl (including the neutrality-preserving Cl^-^), which may aid the retention of water in the vein and perhaps prevent embolism. This would be similar to a report that in short-term assays on tree stems, increasing K^+^ concentration in the xylem perfusate increased radial water hydraulic conductance and has been correlated to enhanced short-term water refilling in embolism repair (Trifilò et al., 2014).

Importantly, in our experiments, ABA or BL regulation of the K_leaf_ occurred independent of stomata regulation, with the BSCs acting as a water valve *in series* with the stomata (Shatil-Cohen et al., 2011; Grunwald et al., 2022). Since both aquaporins (underlying P_f_ and therefore K_leaf_) and ion transporters (including K^+^ channels which we showed to interact with the plant water management) are regulated by pH_EXT_ and pH_CYT_, the ABA-induced alkalization within and outside the BSCs suggests a mechanism whereby drought (mediated by ABA) can affect the BSCs function as a selective barrier to water radial transport into the leaves. Such a mechanism – modifying the BScs’ selective barrier function by altering the pH in the membrane vicinity – could be in operation also under other stresses signaled by ABA, such as cold or salinity. Consequently, our results highlight both proton pumps (AHA2 and VHA) in the BSCs as attractive targets for future designs of plants with stress-tolerance strategies, aiming to improve plants survivability under drought.

## ACKNOWLEDGEMENT

This research was supported by ISF (the Israel Science Foundation, Grant No. 1312/12 to NM and grant No. 1043/20 to MM) and the Ministry of Agriculture, Israel (the Office of the Chief Scientist, grant No. 12-01-0007 to MM&NM). The authors are grateful to Drs. M. Krebs & K. Schumaker for their gift of the double VHA mutant, vha-a2 vha-a3, to Dr. Dizza Bursztyn of the Hebrew University of Jerusalem for advice on the statistics, to Dr. Noa Wigoda – now at the Weizmann Inst of Science, Rehovot, Israel – for occational support in the early stages of the electrophysiology, to Mr. Eyal Erez for the leaf venation data, to Mr. Amir Mayo for the lab light measurements.

## AUTHOR CONTRIBUTIONS

TTS planned and carried out the experiments, MBAR participated in the patch-clamp experiments execution and analyses, YG planned and performed the detached leaf experiments, AD generated SCR:GFP labels for the BSCs and performed the detached leaf experiments, AY generated and contributed the SCR:abi1-1 mutant plants, TTS and VS performed the experiments on whole plants, NM and MM conceived and supervised the experiments, and NM, MM and TTS wrote the paper.

## MATERIALS AND METHODS

### Plant Material

#### Genotypes

We used two accessions of wild type (WT) *Arabidopsis thaliana*: Landsberg *erecta* (Ler) and Columbia (Col), and a few mutants and transformants: *abi1-1* (ABA Insensitive 1-1), the whole plant knockout of a PP2C (*ABI1,* At4G26080.1, in Ler background; Koornneef et al., 1984; Leung et al., 1997; obtained from TAIR, stock CS22); *SCR*:*abi1-1* (Col) plants, i.e., WT(Col) plants transformed with the *abi1-1* gene under the BSCs-directing Scarecrow (*SCR*) promoter (Wysocka-Diller et al., 2000; and see below); *aha2-4* (Col) (SALK_ 082786), a whole-plant T-DNA insertion-knockdown of the H^+^-ATPase, *AHA2* (Haruta et al., 2010; Haruta and Sussman, 2012), obtained from the Arabidopsis Biological Resource Center (Ohio State University); *SCR:AHA2*, *aha2-4* plants complemented with the *AHA2* gene under the BSCs-directed *SCR* promoter (Grunwald et al., 2021), and *vha-a2 vha-a3* (Col), the double mutant of the two tonoplast-localized isoforms of the membrane-integral subunits of VHA, VHA-a2 and VHA-a3 (Krebs, M., Beyhl, D., Görlich, E., Al-Rasheid, K. A. S., Marten, I., Stierhof, Y.D., Hedrich, R., and Schumacher, 2010); a kind gift from Dr. Schumaker’s lab).

#### Protoplasts selection criteria

Most single-cell experiments employed GFP-labeled BSCs protoplasts from the following plant genotypes harboring *SCR:GFP*: WT (Ler); (Shatil-Cohen et al., 2011), WT (Col), *aha2-4* (Col) and *vha-a2 vha-a3* (Col). We used plants from progressively higher generation numbers selected for brighter GFP labeling of BSCs (Attia et al., 2020). In several experiments in which we monitored changes of membrane potential (E_M_, see below) we used protoplasts from plants which did not harbor SCR:GFP and selected the BSCs based on their special appearance: a smaller size and lower chloroplasts density as compared to mesophyll cells (an 70%-successful approach in the hands of an experienced experimenter, already validated by Grunwald et al., 2022, their Suppl. Fig. S11). In the present work, the experiments using BSCs selected based on their “special appearance” were as follows: Figs. 1B-1D, Figs. 2D, 2F, 2G.

#### Plant growth

The Arabidopsis plants were grown in a growth chamber as detailed previously (Grunwald et al., 2021).

#### SCR:abi1-1 (Col) constructs and plant transformation

For construct assembly the MultiSite Gateway Three-Fragment Vector Construction Kit (Invitrogen) was used according to the manufacturer’s instructions. Bundle-sheath-ABA-insensitive (*SCR*:*abi1-1*) plants were constructed similarly to *fa* plants of (Negin et al., 2019).

Quantification of *abi1-1* expression in WT, *abi1-1* and *SCR:abi1-1* plants by restriction enzymes was as described by (Negin et al., 2019; Yaaran et al., 2022).

### Fluorescence imaging

#### Fluorescent dyes for the determination of H^+^-Pumps activities

We monitored the pH and membrane potential (E_M_) resulting from the H^+^-pumping using fluorescence imaging of pH- and E_M_-reporting ratiometric fluorescent probes: (a) pH_EXT_, the pH of the xylem perfusion solution (XPS), using FITC-dextran, FITC-D (fluorescein isothiocyanate conjugated to 10 kD dextran, a dual-excitation pH probe, (Grunwald et al., 2021), perfused via the petiole into the xylem of detached leaves (b) pH_CYT_, the BSCs’ cytosolic pH – using SNARF1 (5-(and 6)-carboxy SNARF-1 acetomethyl ester acetate), a dual emission pH probe, preloaded into the isolated protoplasts (Zhang et al., 2001) and (c) the BSCs’ E_M_ – with di-8-ANEPPS (4-(2-[6-(Dioctylamino)-2-naphthalenyl]ethenyl)-1-(3-sulfopropyl)pyridinium inner salt), a dual excitation potentiometric probe (Pucihar et al., 2009; Wigoda et al., 2017; Grunwald et al., 2022), also preloaded into the isolated protoplasts.

#### pH_EXT_: detached leaves preparation and imaging

The experiments were performed between 9 AM and 1 PM. In the morning of the experiment, shortly before lights-on in the growth chamber (at 9 AM), several 6-7 week old plants were placed in a nearby dark room at roughly the same temperature. Leaves, approximately 2.5 cm long and 1 cm wide, were excised under green light with a sharp blade at 20 min intervals, placed in Eppendorf vials with unbuffered xylem perfusion solution (XPS, see Solutions) containing 100 μM of FITC-D (Ex1: 450, Ex2: 488 nm, Em: 520 nm), added from a 10 mM stock in DDW, as described by Grunwald et al., 2021, without or with the addition of 10 μM ABA. The vials (wrapped in aluminium foil) were placed in humidity boxes (Grunwald et al., 2021), and returned to the growth room for 1.5 -2 hours in full light. Each leaf was then brought back individually for a microscope slide preparation, which lasted about 2 min, it remained on the microscope stage for about 5 min. and during these 2+5 min it remained under *ambient darkness* (the light level not exceeding 0.025 µmol m^-2^ s^-1^ between 400 and 850 nm, as determined using a spectrometer (Maya2000-Pro with the 400 μm head, from Ocean Optics, Germany, www.oceanoptics.eu).

The veins of detached leaves were imaged using *setup I* (based on an inverted microscope Olympus-IX8, as detailed by (Grunwald et al., 2021). Image capture and image analysis of the intra-xylem pH (pH_EXT_), as well as pH_EXT_ calibration were exactly as already described (ibid.), except an updated conversion relationship (pH_EXT_ = (F_ratio-0.06658) / 0.25029, based on an *in-situ* calibration) was used in the current analyses.

#### pH_EXT_; image analysis

The analysis details were as in (Grunwald et al., 2021, 2022).

#### Protoplasts preparation

The experiments were performed between 10 AM and 4 PM. BSC protoplasts were isolated from 6-8 weeks old *SCR:GFP*-labeled plants in an approximately 30 min-long procedure, described earlier (Shatil-Cohen et al., 2014). Upon the collection of the enzymatically released protoplasts, they were washed three times by a 5-fold dilution into the planned experimental bath soluion (see Solutions), which included 0.5 mM ascorbic acid, and were then kept in an Eppendorf vial at room temperature (22 ± 2 °C) under ambient darkness, as above, until use. Generally, 62 μM ascorbic acid remained present in the bath solutions during the experimental treatments, except after the bath solution was flushed out from the experimental chamber.

#### Protoplasts imaging setup and selection criteria

The protoplasts were imaged in *setup II*, consisting of an inverted epifluorescence microscope (Eclipse Ti-S, Nikon, Tokyo, Japan) coupled to an IXON Ultra 888 camera (Andor, UK) via an OptoSplit device (Cairn research, UK). A Nikon 40X objective (Plan Fluor 40x/ 0.75, WD 0.72 mm, DIC) was used for protoplast viewing and imaging. The fluorescence excitation beam was delivered by a xenon lamp monochromator (Polychrome II, Till Photonics, Munich, Germany), under the control of IW6.1 (Imaging Workbench 6.1) software (Indec Biosystems, Santa Clara, CA). All excitation and emission filters were from Chroma Technology Corp. (Bellows Falls, VT, USA).

An individual, perfectly round BSC (diameter of 25-32.5 µm) was selected for imaging based on its GFP fluorescence (Ex.: 490/15 nm, Em.: dichroic mirror: 515 nm and band-pass barrier filter: 525/50 nm), or based on their special appearance, as detailed above.

#### pH_CYT_ using SNARF1; protoplasts treatments

400 μL of bath solution which included 50 μL of protoplasts suspension, SNARF1-AM (5 μM final conc., see Solutions) and pluronic acid (0.05 % final conc., see Solutions), premixed in an Eppendorf vial, were added to the experimental chamber and incubated for 20 min at RT. While the cells settled and stuck to the chamber glass bottom, the membrane-permeant SNARF-AM was digested by cytosolic hydrolases releasing SNARF1, the membrane impermeant free fluorescent ionic form of the dye which largely remained trapped in the cytosol – the most alkaline cellular compartment – for the duration of the recording. At min 10 into SNARF1-AM incubation, red (RL), or blue and red (BL+RL) illumination was aplied for 10 min (from a home-made illuminator of crystal clear LEDs from Tal-Mir electronics, Israel), consisting of RL, 660 nm, roughly 195 μmol m^-2^ s^-1^ and BL, 450 nm, roughly 25 µmol m^-2^ s^-1^ at the protoplast level. During the 10^th^ minute of illumination, the cells were flushed to to replace the incubation solution with SNARF-AM and pluronic acid with a bath solution without or with 3 µM ABA (the flush volume, ≥ 5 mL, was more than 10X the chamber volume of ∼400 μL). Thereafter, the RL / RL+BL illuminator was switched off.

In experiments with **vanadate** (Figs. 3B, 4A and 4B), WT (Col) BSCs protoplasts were incubated in the dark in an Eppendorf tube for 20 min in the standard bath solution (total volume 400 μL) with vanadate (1 mM) and then, for 20 more min also with SNARF1 (5 μM+ 0.05% pluronic acid). Then, the mix with the protoplasts was transferred to the experimental chamber and illuminated with RL+BL (unless otherwise indicated). Vanadate was included also in the flush solution. BAPTA-AM, BAPTA-K_4_, La^3+^ were added along wih SNARF1-AM and flushed away with it.

#### pH_CYT_; imaging using SNARF1

The selected protoplast was focused at its largest diameter under green-filtered phase contrast illumination (‘transmitted light’) and its image was recorded. Immediately thereafter, the SNARF1 within the protoplasts was excited by 550/15 nm 50 ms long light pulse, the emitted fluorescence was deflected by a 570 nm dichroic mirror (T570 LPXR) into the OptoSplit device, further filtered via a 575 nm long-pass filter (ET575LP), then split by a 612 nm dichroic mirror (T612 LPXR-UF2) into two beams, filtered in parallel: via a 640/20 nm band-pass filter (ET640/20M) and via a 585/20 nm band-pass filter (ET585/20M), and the two images were recorded simultaneously on two separate halves of the camera CCD sensor. Then, a next BSC cell was selected in the same chamber, and so on, until 15 min elapsed since the bath flush with the SNARF1-AM-free solution (±ABA). A next batch of cells was then transferred to a clean chamber for further imaging. Experiment-specific deviations from this procedure are detailed below.

#### pH_CYT_; image analysis

The emitted fluorescence intensities in the recorded images were evaluated using FIJI (Abràmoff et al., 2004; Schindelin, J., Arganda-Carreras, I., Frise, E., Kaynig, V., Longair, M., Pietzsch, T., Preibisch, S., Rueden, C., Saalfeld, S., Schmid, B., Tinevez, J.Y., White, Daniel J., Hartenstein, V., Eliceiri, K., Tomancak, P., and Cardona, 2012). The measure of the cytosolic pH was the ratio between the fluorescence intensity at 640 nm which *in*creases with pH, and the fluorescence intensity at 585 nm which *de*creases with increasing pH (Zhang et al., 2001) each corrected first for the corresponding average background fluorescence. To minimize possible errors due to a possibly imperfect overlap of the two recorded images in each pair, the ratiometric analysis was performed on the *averaged* intensities of all the pixels in the image which passed the selection criteria. We applied the automatic thresholding of FIJI to the usually sufficiently bright image at 585 nm and we excluded manually those image regions which were selected automatically but departed obviously from the circumference ring, as well as regions which appeared saturated (the selected areas are enclosed within yellow lines in the Suppl. Fig. S4A).

#### pH_CYT_ calibration using SNARF1

To calibrate the SNARF1 emission ratio values vs cytosolic pH (pH_CYT_) values, we used SNARF1-loaded WT (Ler) protoplasts in solutions containing either 200 mM K^+^ (as in Suppl. Fig. S4B) or 250 mM K^+^ (as in Suppl. Fig. S4C) (see Solutions below). The incubation with SNARF1 and the imaging were conducted as described above except (a) there was no BL+RL illumination, (b) the flushed-in bath solution contained 5 µM nigericin (an H^+^/K^+^ exchanger) and was bufferred to different pHs (pH_EXT_; see Solutions) and (c) the protoplasts were imaged within the first 3-5 min from the nigericin flush-in, before it reached the endomembranes and equilibrated the cytosol with other compartments.

Assuming that an equilibrium is attained within a few minutes across the plasma membrane (Thomas et al., 1979), proton concentrations ratio across the plasma membrane becomes equal to the K^+^ concentrations ratio:

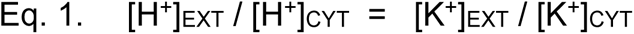

and the internal pH (pH_CYT_) becomes equal to the external pH (pH_EXT_) when [K^+^]_EXT_ = [K^+^]_CYT_.

*Potential error sources of SNARF1 calibration:* Because the [K^+^]_CYT_ in BSCs in our experiments is not known, the resulting deviation of pH_CYT_ from the imposed pH_EXT_ value is also unknown. Thus, if [K^+^]_EXT_ that we use > the true [K^+^]_CYT_, then the *true* pH_CYT_ that we impose > pH_EXT_ which we *think* that we impose. Thus, for example (based on a simulation using Eq. 1), if we use a calibration solution with 200 mM K^+^ at pH 6.5 (i.e., [K^+^]_EXT_ = 200 mM, pH_EXT_ = 6.5), while the true (but unknown) [K^+^]_CYT_ = 400 mM, the resulting (also unknown) pH_CYT_ will be 6.2, rather than 6.5 that we record, but if the the true [K^+^]_CYT_ = 100 mM, the resulting true pH_CYT_ will be 6.8, rather than 6.5 that we record, also unbeknown to us. These potential errors should affect mainly the *absolute* pH values, but the relative pH changes still remain valid.

Moreover, inexplicably, WT (Col) Arabidopsis were recalcitrant to the calibration with 5 µM nigericin, which was therefore performed only on WT (Ler) protoplasts and we assumed its validity also for the other genotypes.

### pHCYT: Specific experimental protocols

*Experiment with SNARF1 and Vanadate* (Fig. 5B). 100 uL of protoplasts suspension (in the bath solution) was incubated in an Eppendof tube with 1 mM vanadate for 30 min. 50 uL of this mixture was then resuspended in an Eppendorf tube in 400 uL of bath solution with 5 uM SNARF1, 0.05% pluronic acid and 1 mM vanadate, and the mix was transferred for 20 min into the experimental chamber, to be illuminated during the 2^nd^ 10 min of the incubation. The flushed-in bath solution contained 1 mM vanadate with or without 3 uM ABA. Cells were then imaged within the subsequent 15 min.

*Experiment with SNARF1 and bafilomycin A1* (Fig. 5C) was conducted as the “standard” SNARF1 experiment, except 100 nM bafilomycin A1 was included in the incubation solution in addition to SNARF1 and pluronic acid. The flushed-in solution was only the bath solution ±3 μM ABA. Cells were then imaged within the subsequent 15 min.

*Experiments with SNARF1 and BAPTA-AM and vanadate* (Figs.4A, 4B). 100 μL of protoplasts suspension (in the bath solution) was incubated in an Eppendof tube with 1 mM vanadate for 20 min. 50 uL of this mixture was then resuspended in another tube in 400 μL of bath solution with 5 μM SNARF-1, 0.05% pluronic acid and 1 mM vanadate without or with 10 μM BAPTA-AM. After another 20 min, the 400 μL mix was transferred to the chamber and illuminated with either RL or RL+BL for 10 min. During the last 1 min of illumination, the cells were flushed with bath solution with 1 mM vanadate with or withour 3 μM ABA. Cells illuminated with RL were then imaged for 5 min (to limit the number of the brief exposures to BL excitation light used for seeking the GFP-labeled BSCs) and cells illuminated with RL+BL were imaged for 15 min.

*Experiment with SNARF1 and BAPTA-AM without vanadate in the dark* (Fig. 4f) was conducted as above, except in the absence of vanadate. The cells remained in the dark throughout their incubation in the tube and in the experimental chamber, and during the flush-in of the bath solution with or without 3 μM ABA. The cells were imaged only for 5 min (to limit the GFP-seeking BL exposures).

*Experiments with SNARF1 and BAPTA-K_4_ and LaCl_3_ in the dark* (Figs. 4G, 4H). Bath solution in these two experiments contained 0.4 mM CaCl_2_ instead of the standard 1 mM. Protoplasts were incubated in this bath solution in an Eppendorf tube with 5 μM SNARF-1, 0.05% pluronic acid and either with 5 mM KCl or 1 mM KCl+1 mM BAPTA-K_4_ (Fig. 4G), or with 5 mM KCl ±50 μM LaCl_3_ (Fig. 4H). Following 10 min incubation in the tube in the dark and 10 min in the experimental chamber in the dark, the cells were flushed with the appropriate bath solution with or without 3 μM ABA, removing all additives except BAPTA-K_4._ The cells were then imaged for 5 min.

*Experiment with SNARF1 and DMSO (a control for the bafilomycin experiment,* Suppl. Fig. S5). 130 mM DMSO was included in the incubation solution in addition to SNARF1 and pluronic acid, and also in the flushed-in solution with ±3 μM ABA. Cells were then imaged within the subsequent 15 min.

### EM: protoplasts imaging

The protoplasts were imaged in the Nikon-microscope-based *setup II*, using Di-8-ANEPPS, as described earlier by Grunwald et al., 2022, with a few changes. A 50 μL drop with leaf protoplasts was resuspended in 400 μL isotonic bath solution containing 30 μM ANEPPS and 0.05% pluronic acid, without or with 3 μM ABA, and the mix was gently added to the chamber, filling it completely. Then, blue and red (BL+RL) light was turned on for 10 min, then turned off. A BSC protoplast, selected based on its GFP fluorescence or based on its special appearance (see protoplasts selection criteria above), was focused at its largest diameter and its transmitted-light image was recorded as above; then it was exposed to a pair of consecutive 3 ms-apart, 50 ms-long excitation pulses, of 438 nm, then 531 nm. The di-8-ANEPPS fluorescence emitted from the membrane (the dye which remained dissolved in the bath did not fluoresce) was filtered via a dichroic mirror of 570 nm and 585/20 nm emission band-pass filter and the pair of the resulting *consecutive* images was recorded by the camera. A next BSC cell was then selected in the same chamber, and so on, for about 5 min, until a total of 15 min elapsed since di-8-ANEPPS (±ABA) addition. A next batch of cells was then transferred to a clean chamber for further imaging.

### EM: Specific experimental protocols

*Experiment with Di-8 ANEPPS and BAPTA-K_4_ and LaCl_3_* (Figs. 1B, 1C, respectively). In both, the modified standard bath solution contained 0.4 mM CaCl_2_ (instead of 1 mM). Additionally, in the experiments of Fig. 1B the bath solution contained either 5 mM KCl (as in the standard solution) or 1 mM BAPTA-K_4_ +1 mM KCl (preserving the K^+^ concentration). After premixing a given bath solution in an Eppendorf tube with 30 µM Di-8 ANEPPS and 0.05 % pluronic acid and protoplasts, the mix was added to the experimental chamber and incubated for 10 min in the dark or under RL or RL+BL. Subsequently, the cells were imaged for 5 min.

*Experiment with Di-8 ANEPPS and BAPTA-AM* (Fig. 2C). A suspension of protoplasts in the standard bath solution premixed with 30 µM Di-8 ANEPPS, 0.05% pluronic acid ±10 µM BAPTA-AM was added to the experimental chamber and incubated for 10 min under RL or RL+BL and the cells were then imaged for 5 min.

*Experiment with Di-8 ANEPPS and bafilomycin* (Suppl. Fig. S2C, S2D) Protoplasts were incubated in an Eppendorf tube for 10 min in the standard bath solution with 30 µM Di-8 ANEPPS, 0.05% pluronic acid and 100 nM bafilomycin A1, then transferred to the experimental chamber and incubated for 10 more min under RL+BL. During the 10^th^ min, a bath solution ± 3 µM ABA (without other additives) was flushed in and the cells were then imaged between 0 - 15 min.

***Image analysis and fluorescence ratio calibration of di-8-ANEPPS*** were as detailed earlier (Grunwald et al., 2022). Briefly, in two, several-months-apart experiments, we obtained a positive relationship between the membrane potential, E_M_, imposed using patch-clamp and the mean fluorescence ratio (F-ratio) of the cell. Importantly, while this F-ratio vs. E_M_ conversion could not be applied quantitatively to the data, due to the variability of the range of the F-ratio values between experiments, we deem it a reliable and sufficient indication of the relative *direction* of the treatment-induced E_M_ changes: depolarization or hyperpolarization.

### Determination of the leaf hydraulic conductivity (K_leaf_)

6-8 weeks old Arabidopsis plant leaves were excised predawn and were put in a vial with their petioles dipped in AXS (artificial xylem solution, see Solutions) with or without 10 mM BAPTA-AM ±10, or with μM ABA and incubated for 1-2.5 hours under light in closed humidity boxes (to maintain humidity). After 1 hour, the box was opened and allowed for equalibration with ambiant humiditiy for 10-20 minutes. Transpiration from each leaf and its water potential were measured using Li-600 porometer (LI-COR) *and* pressure chamber, respectively as described in Grunwald et al., 2021. K_leaf_ was calculated as described in Grunwald et al., 2021.

### Electrophysiology

***Patch clamp technique*** for determination of K^+^ channel activity in the BSCs. The experiments were performed using a Digidata 3122A interface, Axopatch 1C amplifier and pClamp 10.2 program suite (all three from Molecular Devices, Union City, CA), used both for running the experiment and for analysis. The experiments were done at 22±2°C. Patch-clamp pipettes were pulled from borosilicate glass capillaries on a horizontal P-87 puller (both from Sutter Instruments, Novato, CA; capillaries Cat. No. BF150-86-10) and were "fire-polished’ against a heated Pt-Ir filament, dipped in protamine sulfate (Sigma P-4380, 1% in water) and dried, then filled with ‘pipette solution’ (see ‘Solutions’ below), and the fire-polished pipette tip was coated with wax. A drop of wash solution with protoplasts was added to the 5 mM K^+^ bath solution at a selected pH in the experimental chamber (all the solutions are listed below under ‘Solutions’) and the protoplasts were allowed to settle for 10 minutesSubsequently, a suitable protoplast was selected (based on aits GFP label and a diam. Range of 25-32.5 um). Whole-cell configuration was usually attained spontaneously upon a slight suction on the patch pipette without first obtaining a ‘giga-seal’ (Ward, 2014). Usually, within a few minutes at a holding potential of 0 mV, the baseline current decreased to a null or close to null and stabilized (0 mV is a nominal value; after a ‘Liquid Junction potential’ (LJP) correction (see below) the holding potential was -23 mV). The pipette resistance was monitored by a repetitive testing pulse of 10 mV. The capacitive transients were cancelled using the compensatory circuit of the patch-clamp amplifier. All experiments were performed in a ‘voltage-clamp’ mode. The error in the voltage clamping of the whole-cell membrane, largely due to the series resistance (Rs) of the patch pipette, was compensated at approximately 80% by analog circuitry of the Axopatch 1C amplifier. The voltage error due to the residual Rs was less than 10% of the membrane potential. Currents were filtered at 1000 Hz and sampled usually at 2 kHz. Membrane currents were elicited from the protoplast membrane by repeated “sweeps”, each consisting of a 5 s long square voltage pulse (‘pre-pulse’) decreasing from sweep to sweep at -20 mV steps, from +140 to -180 mV; each square pre-pulse was followed by a brief 4.5 ms-long ramp from +70 to -70 mV (all the voltage values mentioned here are nominal, before the LJP correction, but see below), referred in the Results section as “E_rev_ -testing ramp”.

The brevity of the ramps ensured that the membrane conductance during the ramp reflected the conductance prevailing during the moment just preceding the ramp. The total sweep duration did not exceed 5 s, and the inter-sweep interval was 30s.

#### Patch-clamp data analysis: *LJP correction*

Based on the composition of the pipette and bath solutions, the values of the membrane potential (E_M_) were corrected for the Liquid-Junction potential (LJP) calculated using the pClamp 10.2 suite calculator (and added to all values of E_M_). Thus for the pair of “internal” (pipette) and “external” (bath) solutions, LJP was -23 mV. All the values of E_M_ mentioned henceforth are already corrected for LJP.

#### Reversal potential - channel selectivity

The E_rev_ -testing ramp served to generate linear current-voltage (I-E_M_) plots with different slopes (G_net_), each corresponding to a different channel opening /or closing prepulse, the intersection of which yielded the value of E_rev_, according to the Ohmic relation, Eq. 1:

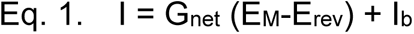

where G_net_ is the net voltage-dependent membrane conductance contributed by the voltage-dependent open channels; E_M_ is the membrane potential and E_rev_ is the membrane potential at which 0 (zero) current flows through presumably a single type of open channels; I_b_ is the basal current which flows through pathways other than through the voltage dependent channels (here: “leak”). For an ideally ion-selective channel, E_rev_ = Nernst potential (an equilibrium potential) for that ion.

#### Specific Membrane Conductance and Bolzmann fitting

The slopes of the (I-E_M_) plots generated by the E_rev_ -testing ramps served for the determination of the specific conductance G’ related to the E_M_ during the prepulse. These G’ -E_M_ relationships served to characterize the channel type, especially when fitted with the Boltzmann relationship (Suppl. Protocols S4, in Wigoda et al., 2017) to yield the best-fit characteristic parameters G’_max_net_ and E_1/2_ (ibid.).

### A Physiological-phenotyping platform in a greenhouse

Using the functional telemetric platform comprised of weighing lysimeters, soil and atmosphere sensors (Plantarray, PA 3.0, PlantDitech Ltd., Yavne, Israel; https://www.plant-ditech.com/ ), plant growth and water balance were continuously monitored through controlled tracking and measurement of the transpiration and biomass gain of each quadruplet of Arabidopsis plants in a pot throughout the growing period. In parallel, also the soil water and atmospheric conditions around the plants were monitored as described in (Dalal et al., 2020).

#### Drought treatment

The plants were grown during two winter seasons, in the phenotyping greenhouse (14 Feb – 21 March, 2019 and 2 Jan – 3 Feb, 2022).

During the 1^st^ season irrigation was withdrawn starting on the night between the 29^th^ and the 30^th^ of Feb 2022, and lasted till the experiment end.

During the 2^nd^ season irrigation was withdrawn starting on the night between the 13^th^ and the 14^th^ of Jan 2022, and was resumed on the night between the 26^th^ and the 27^th^ of Jan 2022, followed by 8 days of recovery.

### Data presentation and Statistics

Individual data, from at least three independent experiments, were usually presented as symbols in box plots using Origin (OriginProVersion 2022. OriginLab Corporation, Northampton, MA, USA). The box in a graph indicates 25 - 75 % of the data range, the red line – the median, “x” – the mean, and the whiskers delimit 3 times the ‘inter-quartile range’ (IQR). Outliers were defined as being outside this three x IQR and were excluded from the statistics. Different treatments / genotypes were compared using the 1-way ANOVA Tukey-HSD test (as implemented by JMP (JMPPro, Version 16.0.0 (512257), SAS Institute Inc., Cary, NC, USA, 1989–2021) or Origin) and labeled with different letters whenever statistically different at P<0.05. When only two groups were compared, we applied the Excel’s Student’s 2-tailed, equal variance t-test (Microsoft Excel, ver. 2108) using the same criterion of P<0.05 for a significant difference. A single-tail t-test was performed only on data of suppl. Fig. S3C.

### Solutions and reagents

***XPS, xylem perfusion solution*** (in mM): 10 KNO_3_, 1 KCl, 0.3 CaCl_2_ and 20 D-sorbitol. Upon preparation, when unbuffered, the pH of this solution was 5.6 - 5.8 and its osmolarity was approx. 23 mOsm.

***Bath solution for SNARF1 (pH_CYT_) calibration (in mM)***: a) *- 200 mM K^+^*, as in Suppl. Fig. S4B (in mM): 200 K-Gluconate, 1 CaCl_2_, 4 MgCl_2_, 10 HEPES, different pHs (pH 6.0, 6.5,7,7.5,8 and 8.5) were adjusted using 1 M NMG, osmolality: 435 mOsm, adjusted w/ D-sorbitol; b) *- 250 mM K^+^*, as in Suppl. Fig.S4C (in mM): 250 K-Gluconate, 1 CaCl_2_, 4 MgCl_2_, 10 HEPES, different pHs (pH 6.5,7,7.5,8 and 8.5) were adjusted using 1 M D-NMG base, osmolality: 456 mOsm.

### Bath solutions for monitoring the protoplasts’ pH_CYT_

The standard solution contained (in mM): 5 KCl, 1 CaCl_2_, 4 MgCl_2_, 10 MES; pH 5.6 (adjusted with 0.5 mM D-NMG base), osmolality: 435 mOsm, adjusted w/ D-sorbitol. In experiments with BAPTA-K_4_, only 0.4 mM CaCl_2_ was used for all treatments; in the presence of 1 mM BAPTA-K_4_, the free [Ca^2+^]_EXT_ was 0.6 μM, as calculated using ‘Extended Maxchelator’ (see details below); Alternatively, at pH_EXT_ 7.5, with 10 mM HEPES (adjusted with 1.6 mM D-NMG base), the free [Ca^2+^]_EXT_ was 0.04 μM, as calculated using the ‘Extended Maxchelator’ (see details below).

### Bath solution for monitoring the protoplasts’ E_M_

The standard solution was as above. The solutions used in experiments with 1 mM BAPTA-K_4_ were also as above.

***Pipette solution for di-8-ANEPPS / E_M_ calibration*** (in mM): 112 K- Gluconate, 28 KCl, 2 MgCl_2_, 2 MgATP (Sigma-Aldrich, Cat. # A6419), 2 BAPTA-K_4_, 1 CaCl_2_ (245 nM free [Ca^2+^]_CYT_, pH 7.5 (10 HEPES, 1.2 D-NMG) base, osmolarity: 480 mOsm (328 Sorbitol).

***Bath solution for patch-clamp experiments*** (in mM): a) at *pH_EXT_ 5.6*, the same as for E_M_ monitoring*, above, b)* at *pH_EXT_ 7*, the same, but with additional 0.5 mM D-NMG base.

***Pipette solution for patch-clamp experiments*** (in mM): the composition was as under “*Pipette solution for di-8-ANEPPS / E_M_ calibration*” above, except for the varying pH (pH_CYT_), added D-NMG and Ca^2+^, and the resulting varying free Ca^2+^ concentration, calculated using ‘Extended Maxchelator’ (Table 1). Throughout the MS, the free [Ca^2+^]_CYT_ of Table 1 are approximated as follows: 127-132 nM – as 100 nM, 557-577 nM as 600 nM.

**TABLE 1:**
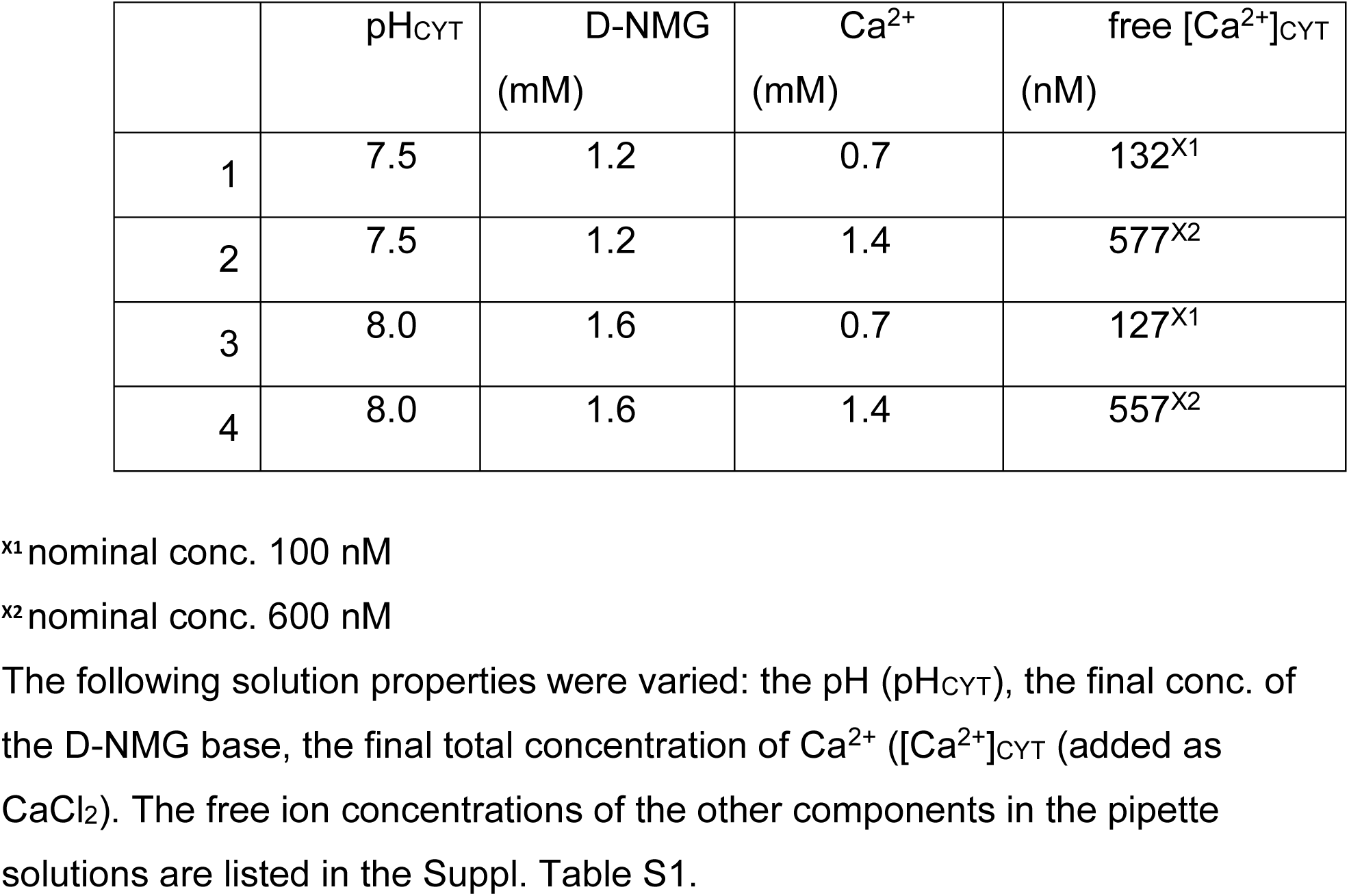
Calculated free Ca^2+^ concentration in the five types of pipette solutions

| | $\text{pH}_{\text{CYT}}$ | D-NMG<br>(mM) | $\text{Ca}^{2+}$<br>(mM) | free $[\text{Ca}^{2+}]_{\text{CYT}}$<br>(nM) |
| --- | --- | --- | --- | --- |
| 1 | 7.5 | 1.2 | 0.7 | $132^{\text{x1}}$ |
| 2 | 7.5 | 1.2 | 1.4 | $577^{\text{x2}}$ |
| 3 | 8.0 | 1.6 | 0.7 | $127^{\text{x1}}$ |
| 4 | 8.0 | 1.6 | 1.4 | $557^{\text{x2}}$ |

<sup>x1</sup> nominal conc. 100 nM
<sup>x2</sup> nominal conc. 600 nM

***ABA:*** (±)-Abscisic acid (Sigma-Aldrich, Cat. # D1049); a 100 mM stock solution in 1 M KOH in DDW was stored in aliquots, protected from light at -20°C. ***FITC-D*:** Fluorescein isothiocyanate–dextran (average mol wt 10,000; Sigma-Aldrich, cat. #: FD10S); a 10 mM stock solution in DDW was stored in aliquots, protected from light, at -20°C.

***Di-8-ANEPPS***: (4-(2-[6-(Dioctylamino)-2-naphthalenyl]ethenyl)-1-(3- sulfopropyl)pyridinium inner salt, Enzo, Farmingdale, NY,USA, cat #: ENZ-52204) a 10 mM stock solution in DMSO (Dimethyl sulfoxide, Sigma-Aldrich, cat. #: D2650) was stored in 10 µL aliquots at -20 °C.

***SNARF1-AM:*** (Carboxy seminaphthorhodafluor-1, acetoxymethyl ester), acetate, Molecular Probes/Thermo Fisher Scientific, Oregon, USA, cat. #: C-1272); 50 µg portions were dissolved in DMSO sequentially, as needed; a 10 mM stock solution was stored in aliquots at -20°C..

***Pluronic acid F-127***: (Molecular Probes/Thermo Fisher Scientific, Oregon, USA, cat. #: P6867), a 20% stock solution in DMSO was stored in 100 µL aliquots at RT.

***Vanadate:*** sodium orthovanadate, Na_3_VO_4_ (BHD Chemicals Ltd., cat. #30194); the stock solution of 200 mM in DDW was depolymerized by boiling as described in the Sigma-Aldrich Product Information sheet for sodium orthovanadate: https://www.sigmaaldrich.com/content/dam/sigmaaldrich/docs/Sigma/Product_Information_aldrich/docs/Sigma/Product_Information_Sheet/1/s6508pis.pdf.

The stock solution was aliquoted and stored at -20 °C.

***Bafilomycin A1*:** a VHA inhibitor (Sigma-Aldrich, cat. #: B-1793); a 0.16 mM stock in DMSO and stored in aliquots, protected from light at -20 °C.

***Nigericin free acid:*** a K^+^/H^+^ ionophore (Molecular Probes/Thermo Fisher Scientific, Oregon, USA, cat. #: N-1495); a 5 mg/mL stock in EtOH (Ethanol 99.5%, HPLC grade, J.T. Baker /Thermo Fisher Scientific, Oregon, USA, cat #: 64-17-5), stored in aliquots at -20°C.

***BAPTA-K_4_:*** 1,2-bis(2-Aminophenoxy)ethane-N,N,N′,N′-tetraacetic acid tetrapotassium salt, Invitrogen/Thermo Fisher Scientific, Oregon, USA, cat. #: B1204); a 200 mM stock solution in H_2_O was stored in 20 μL aliquots at -20 °C.

***BAPTA-AM:*** 1,2-Bis(2-aminophenoxy)ethane-N,N,N’,N’-tetraacetic acid tetrakis(acetoxymethyl ester, MCE, MedChemExpress, Monmouth Junction, NJ, USA, cat. #: HY-100545); a 65 mM stock solution in DMSO was stored in 10 μL aliquots at -20 °C.

***LaCl_3_*:** Lanthanum chloride (Sigma-Aldrich, cat #10009958-8)

***D-NMG base:*** N-Methyl-D-glucamine (Sigma-Aldrich, cat # 66930)

***MaxChelator: Calculation of the free [Ca^2+^] in the presence of a chelator, BAPTA-K_4_***. We used the current version of free software from UCDavis, Extended MaxChelator: https://somapp.ucdmc.ucdavis.edu/pharmacology/bers/maxchelator/webmaxc/webmaxcE.htm, originally by Chris Patton from the University of Stanford.

## SUPPLEMENTAL MATERIALS

### Supplemental figures

FIGURE S1. In light, the hydraulic conductance (K_leaf_) of detached Arabidopsis leaves depends on the concentration of free cytosolic Ca^2+^ ([Ca^2+^] ).

FIGURE S2. ABA-induced depolarization is delayed beyond the first 5 min of ABA exposure and is independent of VHA activity.

FIGURE S3. ABI1, a PP2C (protein phosphatase 2C) mediates the ABA-induced alkalinization of the xylem perfusate in detached leaves

FIGURE S4. Monitoring the cytosolic pH (pH_CYT_) by SNARF1, the fluorescent, dual-emission, ratiometric pH probe.

FIGURE S5. DMSO, at a conc. exceeding those used in our experiments, affects neither the pH_CYT_ of WT (Col) BSCs protoplasts nor their capability to respond to ABA.

FIGURE S6. In light, the hydraulic conductance of detached Arabidopsis leaves (K_leaf_) depends on ABA and on the presence of the K^+^-efflux channel SKOR.

FIGURE S7. The normalized momentary transpiration of greenhouse-grown plants on three representative days during the experiment of Fig. 5C-F.

Figure S8. K^+^ accumulation in fragments of leaf blades excised from the same WT (Col) and *skor* (Col) mutant Arabidopsis plants before and after a two weeks irrigation withholding.

FIGURE S9. Anatomical comparisons between leaf blades of WT (Col) and *skor* (Col).

### Supplemental Tables

TABLE S1. The calculated free concentrations of ions in the patch-clamp pipette solutions listed in Table 1.

### In support of the cell model details: Publications and our experimental data figures

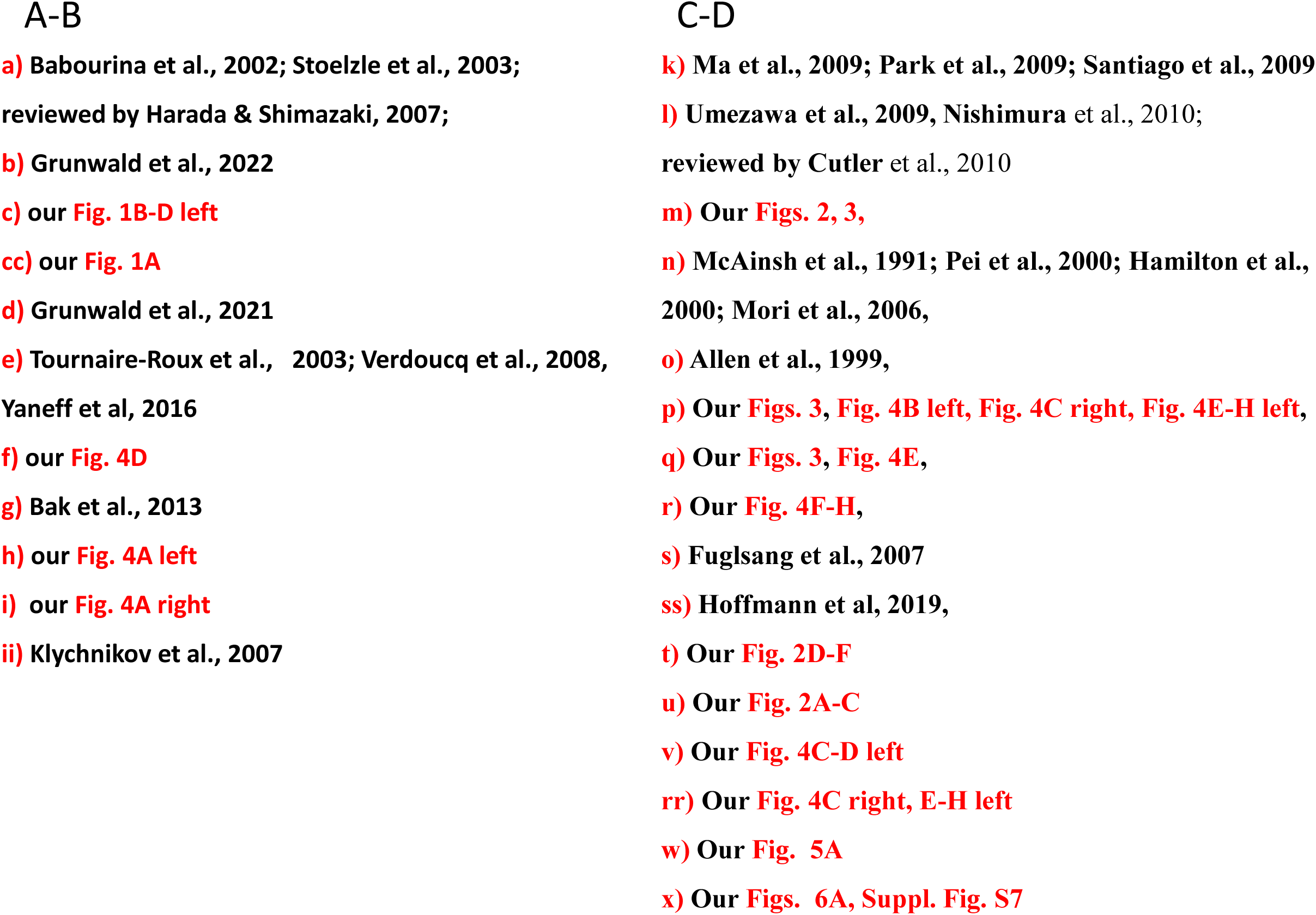

Above-mentioned publications not listed in the main REFERENCES list

