## Supplemental Figures for "A tale of two pumps: Blue light and ABA alter Arabidopsis leaf hydraulics via bundle sheath cells’ H^+^-pumps and channels"

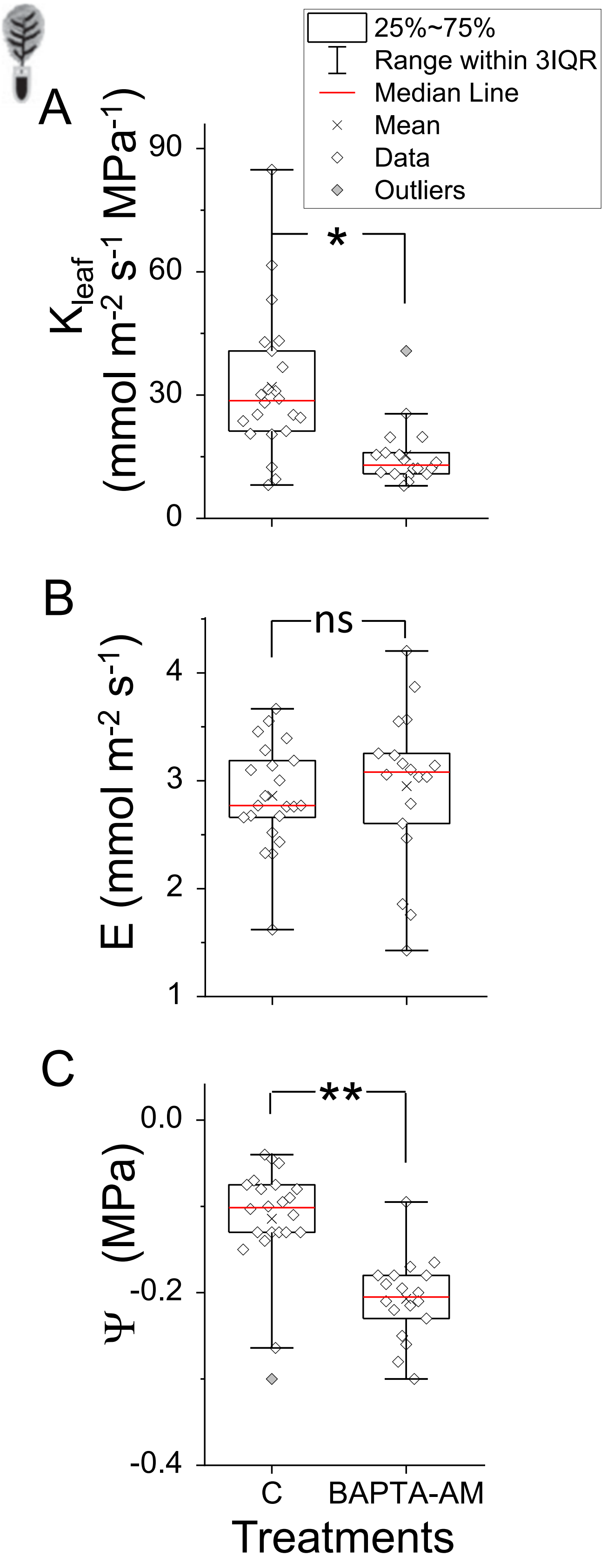


**FIGURE S1. In light, the hydraulic conductance ($K_{leaf}$) of detached Arabidopsis leaves depends on the concentration of free cytosolic $Ca^{2+}$ ($[Ca^{2+}]_{CYT}$).** **A.** The effect of the cytosolic $Ca^{2+}$ chelator, BAPTA-AM (10 µM) on the $K_{leaf}$ of WT (Col) detached leaf. *: a difference from control (C; P = 0.000291, 2-tailed t-test, equal variance; outliers excluded). This part is identical to Fig. 1A. **Inset**, a detached leaf schematics. **B.** The transpiration rate (E). ns: not significant. Note that E is unaffected by the decline of free $[Ca^{2+}]_{CYT}$ – presumably, in the BSCs – suggesting an indifference of stomata conductance to the petiole-introduced $Ca^{2+}$ chelator. **C.** The leaf water potential ($\Psi_{leaf}$). **: a significant difference (P = 9.549E-08, t-test, as in Fig. 1A). The statistics does not include outliers. Note that the decline of $[Ca^{2+}]_{CYT}$ decreased the $\Psi_{leaf}$, indicating decreased water entry into the leaf via the BSCs layer. This is consistent with the BAPTA-AM-induced $K_{leaf}$ decrease being associated mainly with the BAPTA-AM-induced decline in $\Psi_{leaf}$. **Inset:** A detached leaf schematics. *In support of Fig. 1A.*

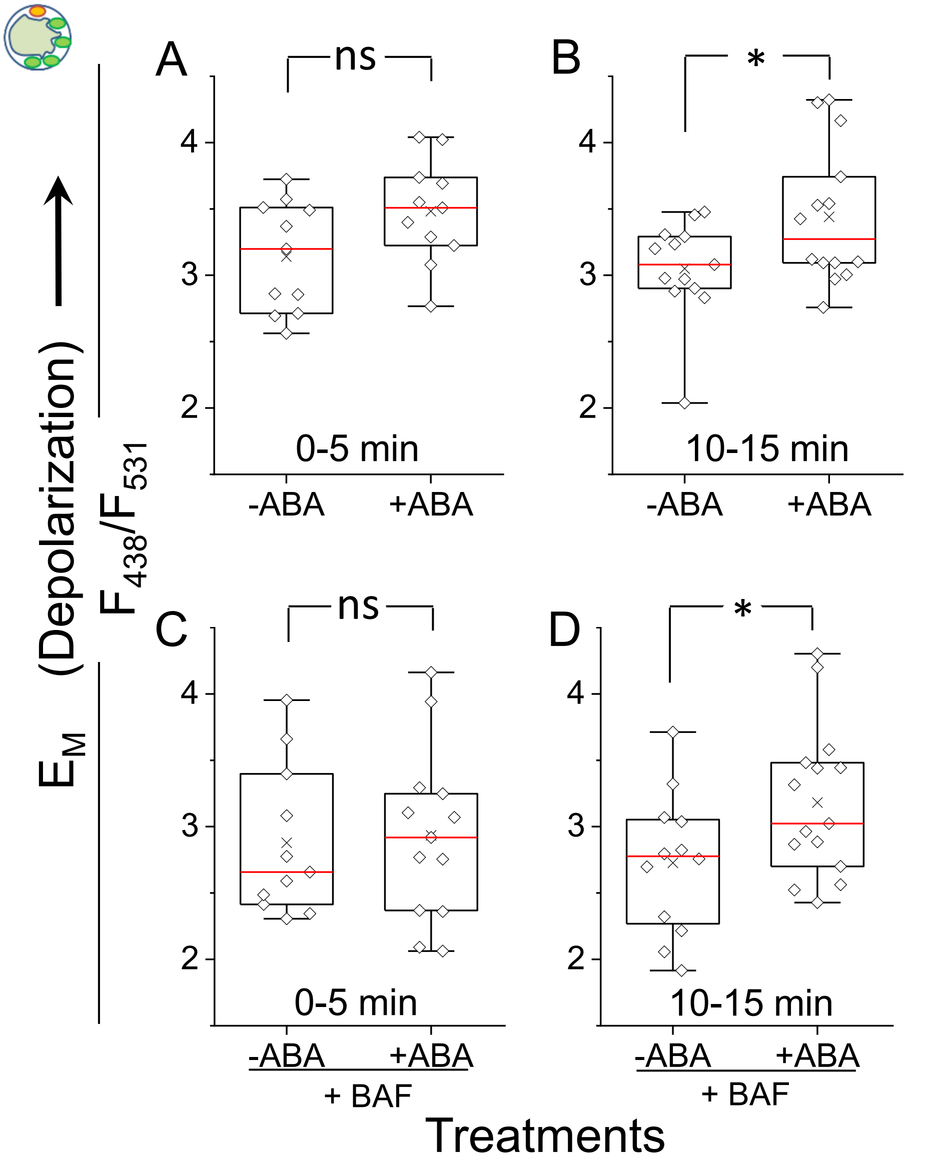


**FIGURE S2. ABA-induced depolarization is delayed beyond the first 5 min of ABA exposure and is independent of VHA activity. A-B.** Membrane potential ($E_M$) of BSCs WT (Col) protoplasts reported by di-8-ANEPPS, as in Fig. 1B, except with the following alterations: (a) the standard solution used in the incubation and in the bath contained additionally 5 mM $KNO_3$, (b) the protoplasts were incubated with di-8-ANEPPS (30 μM) in an Eppendorf tube for 10 min, then in the experimental chamber for an additional 10 min during the RL+BL illumination. During the last min, the same bath solution with $KNO_3$ ± ABA (3 μM) but without other additives was flushed into the experimental chamber and the cells were then imaged for between 0 - 15 min. **A.** Cells imaged within the first 5 min after the exposure to ABA. **B.** Cells imaged within 10-15 min after the exposure to ABA. *: a significant difference (P=0.0345, t-test, as in Fig. 1A). *Supports Figs. 1B - 1D, 2D -2G.* **C-D.** ABA effect on the $E_M$ of WT (Col) BSCs assayed as in A-B, except, during the 20 min incubation with di-8-ANEPPS the protoplasts were exposed also to the VHA-specific inhibitor bafilomycin A1 (BAF, 100 nM), and the standard solution was used in the incubation and in the experimental chamber throughout the experiment. In *support of Fig. 3.* **C.** Cells imaged within the first 5 min after the exposure to ABA. **D.** Cells imaged within 10-15 min after the exposure to ABA. *: a significant difference (P=0.0442, t-test, as in Fig. 1A). Since the ABA-induced depolarization lags appreciably after the fast ABA-induced cytosol alkalinization, it is unlikely that AHA2 inhibition depends on VHA stimulation and cytosol alkalinization. **Inset.** An illuminated BSC protoplast.

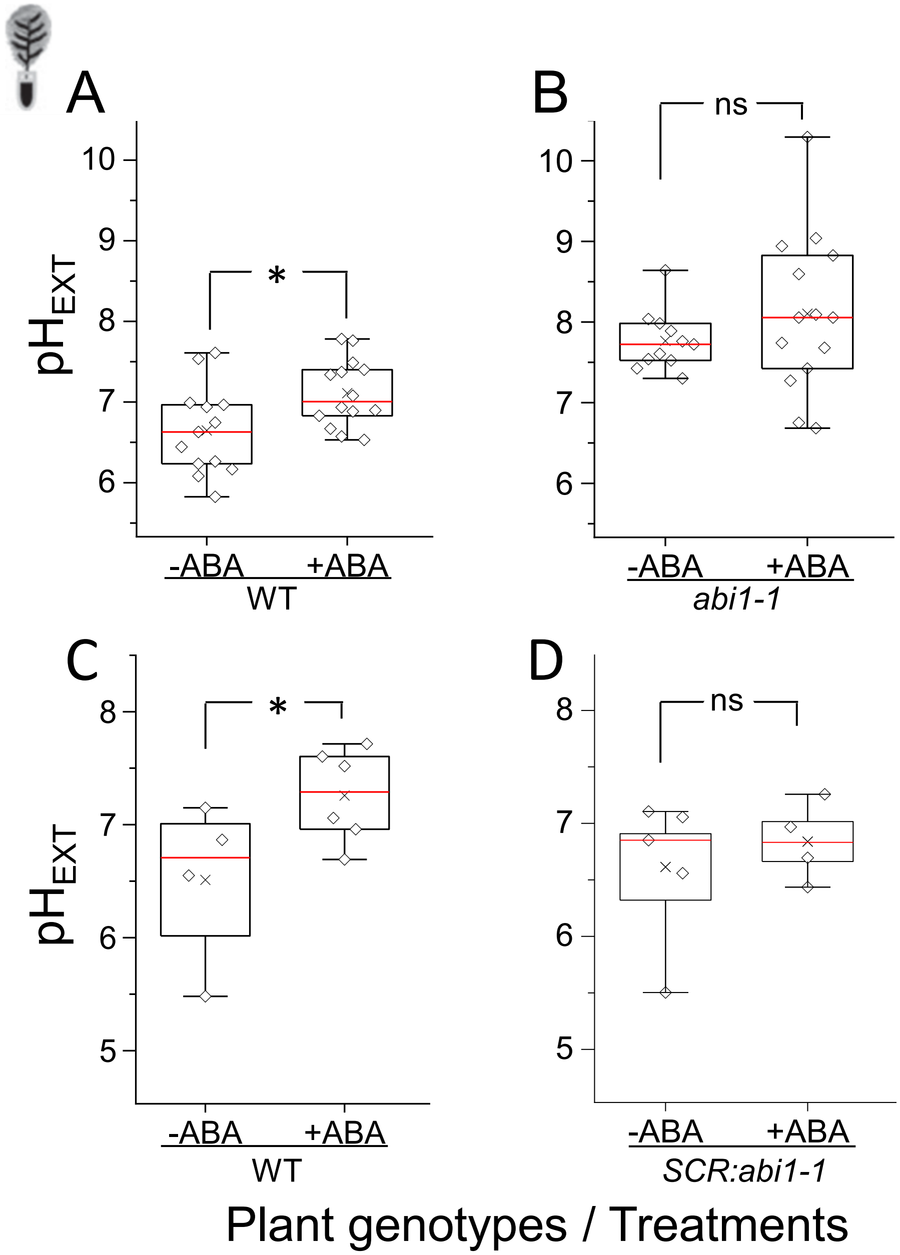


**FIGURE S3. ABI1, a PP2C (protein phosphatase 2C) mediates the ABA-induced alkalinization of the xylem perfusate in detached leaves. A-B.** Data from three independent experiments on detached leaves of WT (Ler). **A.** ABA (10 µM) elevates the pH ($pH_{EXT}$) of the non-buffered xylem perfusion solution (XPS) by 0.3 pH units (from pH 6.7 to 7). *: a significant difference (P=0.02047, t-test, as in Fig. 1A). **B.** ABA has no effect on the $pH_{EXT}$ in the leaves of the *abi1-1* (*ABA insensitive1-1*, Ler) mutant. ns: non-significant difference. **C-D.** Data from two independent experiments on detached leaves of WT (Col), included here due to their confirmatory nature. *Support Fig. 2G.* **C.** ABA alkalinizes the XPS by 0.8 pH units (roughly from pH 6.5 to 7.3). *: a significant difference (P= 0.03484, by a single-tail t-test, equal variance). **D:** ABA has no effect on the $pH_{EXT}$ of WT (Col) Arabidopsis transformed with the mutated *abi1-1* (*ABA insensitive1-1*) gene directed to the bundle sheath cells by the Scarecrow (*SCR*) promoter. Other details as in A. Note the abolishment of ABA effect on $pH_{EXT}$ in the *SCR:abi1-1* plants harboring the *ABA-insensitive* phosphatase ABI1 activity in their BSCs. **Inset.** A detached illuminated leaf.

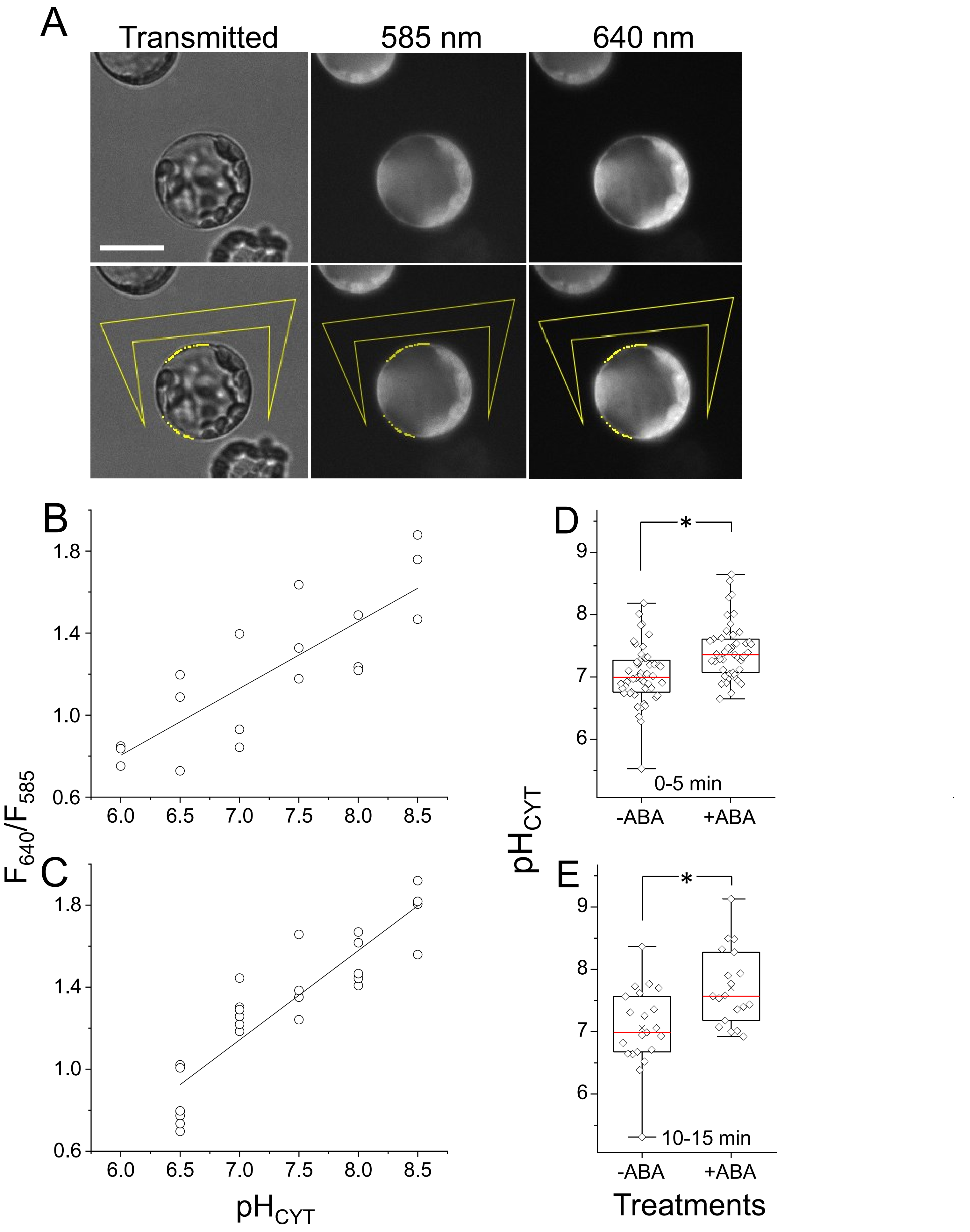


CONT. ON THE NeXT PAGE

Suppl. Fig. S4. Torne-Srivastava et al., 2023

CONT. FROM THE PREV. PAGE

**FIGURE S4. Monitoring the cytosolic pH ($pH_{CYT}$) by SNARF1, the fluorescent, dual-emission, ratiometric pH probe.** **A.** Representative images. Top panel. Transmitted light and fluorescence images of a BSC protoplast (selected based on its GFP fluorescence, not shown) after incubation with the membrane-permeant (-AM) form of SNARF1 and its subsequent washout, recorded at the indicated wavelengths (Ex: 550 nm, $Em_1$:585 nm, $Em_2$: 640 nm). After hydrolysis of the probe by cytosolic hydrolases, the fluorescence emitted from the cytosol of the SNARF1-loaded protoplasts is seen as a bright cell contour in the focal plane. Lower panel: Areas with the most highly fluorescing (but well below saturation) pixels at 585 nm were selected automatically using FIJI and were delimited automatically by yellow lines. Out of these, areas in a ring closest to the cell contour (presumed to represent the cytosol) were accepted manually. The straight lines-delimited area (selected manually) near the protoplasts served for background determination. The selected pixel areas were then superimposed on the corresponding pixels in the 640 nm image and, for orientation, also on the transmitted-light image (Transmitted). The scale bar: 20 µm. A NOTE: The image at 585 nm had its brightness automatically increased by the FIJI analysis program; therefore, in compensation, we tuned it down (by -22% using the "picture format" of PowerPoint) to match that brightness of the unprocessed image above it. **B-C.** *in-situ* calibration of the cytosolic pH in SNARF1-loaded BSC protoplasts performed in the presence of the $H^+/K^+$ exchanger, nigericin (5 µM) and a high $K^+$ conc. in the bath ($[K^+]_{EXT}$) presumed to equal the cytosolic $[K^+]_{CYT}$. **B.** $[K^+]_{EXT}$ = 200 mM. $F_{640}/F_{585}$: the F-ratio, the ratio of the fluorescence emitted from the selected areas at the indicated emission wavelengths vs the pH of the buffered bath solution ($pH_{EXT}$). The symbols are data from individual cells (biological repeats). The fitted line: F-ratio = 0.32564*pH - 1.14968 (Pearson's R = 0.821); for converting the F-ratio values to pH we used: **pH = (F-ratio +1.1497) / 0.3256. C.** An *in-situ* calibration as in B, after a replacement of the excitation source lamp and with $[K^+]_{EXT}$ = 250 mM in the bath. The fitted line: F-ratio = 0.4359*pH - 1.9088 (Pearson's R = 0.904); for the F-ratio conversion to pH we used**: pH = (F-ratio+1.9088) / 0.4359. D-E.** The effect of ABA (3 µM) on $pH_{CYT}$. **D-E.** A collection of $pH_{CYT}$ values reported by SNARF1 from WT (Ler) protoplasts from 13 experiments over a period of several months. **D.** Data from minutes 0-5 of the ABA exposure. Note that ABA alkalinized the cytosol within a few min of its introduction into the experimental chamber. *: a significant difference (P=5.384 $10^{-5}$, t-test, as in Fig. 1A). **E.** As in D, except the data were collected during minutes 10-15 of ABA exposure from a large part of the experiments of D. *: a significant difference (P = 0.02579). *Supports Figs. 3, 4.*

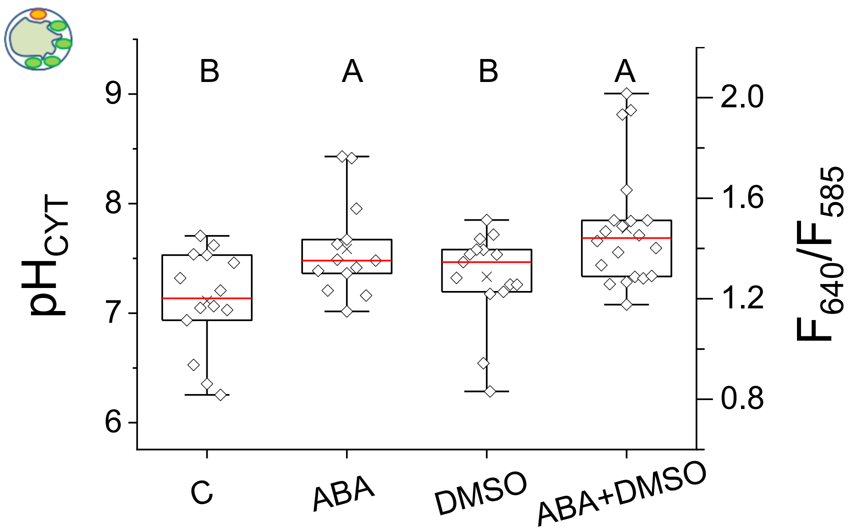


**FIGURE S5. DMSO, at a conc. exceeding those used in our experiments, affects neither the $pH_{CYT}$ of WT (Col) BSCs protoplasts nor their capability to respond to ABA.** The undiluted DMSO concentration is 14.1 M (its density: 1.1 g/cm³ and its molar mass: 78.13 g/mol).

DMSO is the solvent of bafilomycin A1 (at a 1600x final dilution, final conc. 9 mM), BAPTA-AM (at a 6500x final dilution, final conc. 2 mM), di-8 ANEPPS (at a 333x final dilution, final conc. 42 mM), SNARF1-AM (at a 2000x final dilution, final conc. 7 mM) and pluronic acid (at a 400x dilution, final conc. 35 mM). The experiment was conducted as in Fig. 3A left, except DMSO was added, where indicated, at a 160x dilution (final conc. 90 mM), to the incubation solution in addition to DMSO already included in the solutions of SNARF1 and of pluronic acid (together, final DMSO conc. was 132 mM), and also to the flush-in bath solution ("only" 90 mM during the imaging). Different letters indicate statistically different means (at $P<0.05$, ANOVA, Tukey HSD test). Note that $pH_{CYT}$ without or with ABA was not affected by the presence of DMSO. The conversion from Ratio values to pH was done using the calibration in Suppl. Fig. S4C. *In support of Figs. 1-4*

**FIGURE S6. In light, the hydraulic conductance of detached Arabidopsis leaves ($K_{leaf}$) depends on ABA and on the presence of the $K^+$-efflux channel SKOR. A.** Artificial xylem sap (AXS) alone (C or -ABA), or +ABA (10 μM) was imbibed via a petiole into the leaves of WT (Col) or *skor* (Col) mutant plants and one to 2.5 hs later the transpiration rate (E) and the leaf water potential ($\Psi_{leaf}$) were determined), and $K_{leaf}$ was calculated (as in Shatil-Cohen et al., 2011). Other details as in Fig. S1. The data of C are from Fig. S1. Different letters indicate statistically different means (ANOVA, Tukey-HSD test, P<0.05). **: a significant difference (at P = 0.000565, two-tailed t-test, equal variances, or P = 0.000594 by the same t-test on log10 of $K_{leaf}$). The statistics does not include outliers. **B.** The transpiration rate (E) declines sharply under ABA treatment, indifferent to the genotype, indicating the same response in stomata conductance. Other details as in A. **C.** The leaf water potential ($\Psi_{leaf}$) declines under ABA treatment, but less so in the mutant *skor* than in WT. *: a significant difference (P = 0.01214, t-test, as above). Other details as in A. Note that the $\mathbf{K_{leaf}}$ in the ABA-treated *skor* plants declined less than the $\mathbf{K_{leaf}}$ in the ABA-treated WT leaf, associated mainly with the much less pronounced decrease of the $\mathbf{\Psi_{leaf}}$ in the *skor* leaf. *In support of Fig. 5B.*

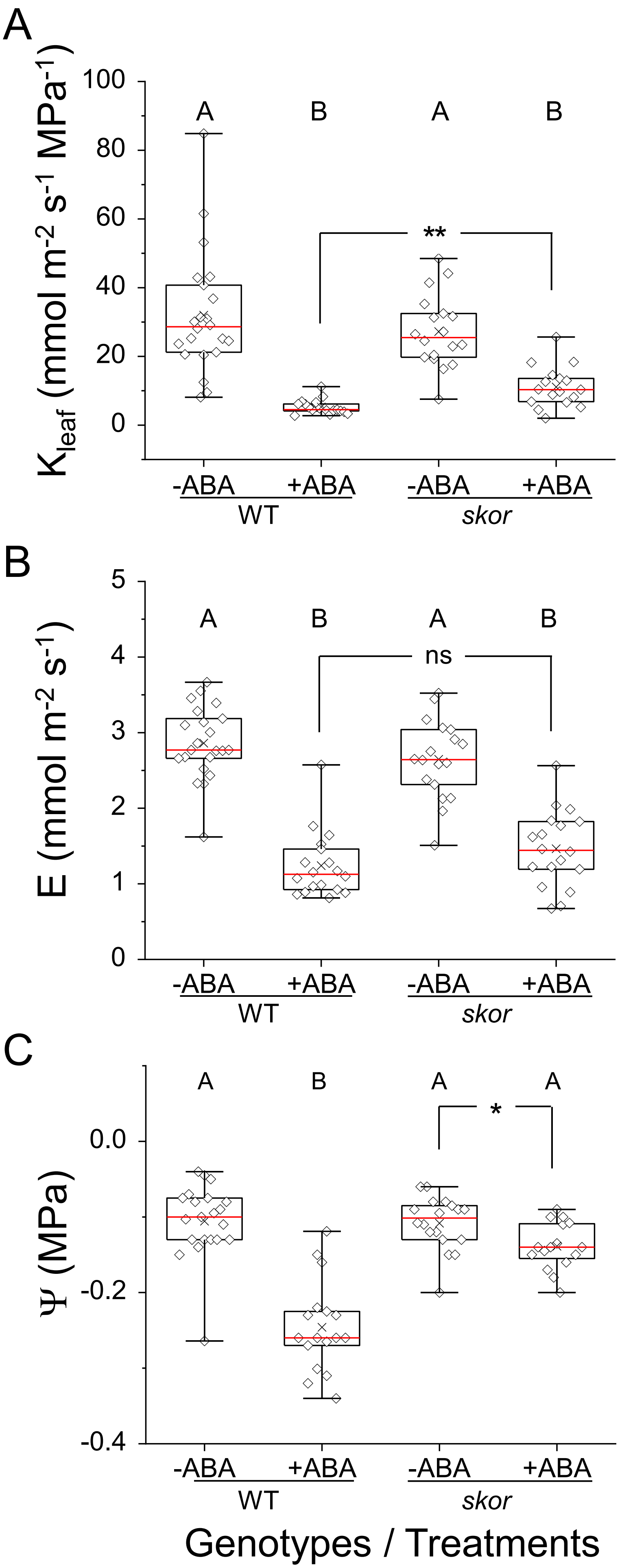


*Suppl. Fig. S6. Torne-Srivastava et al., 2023*

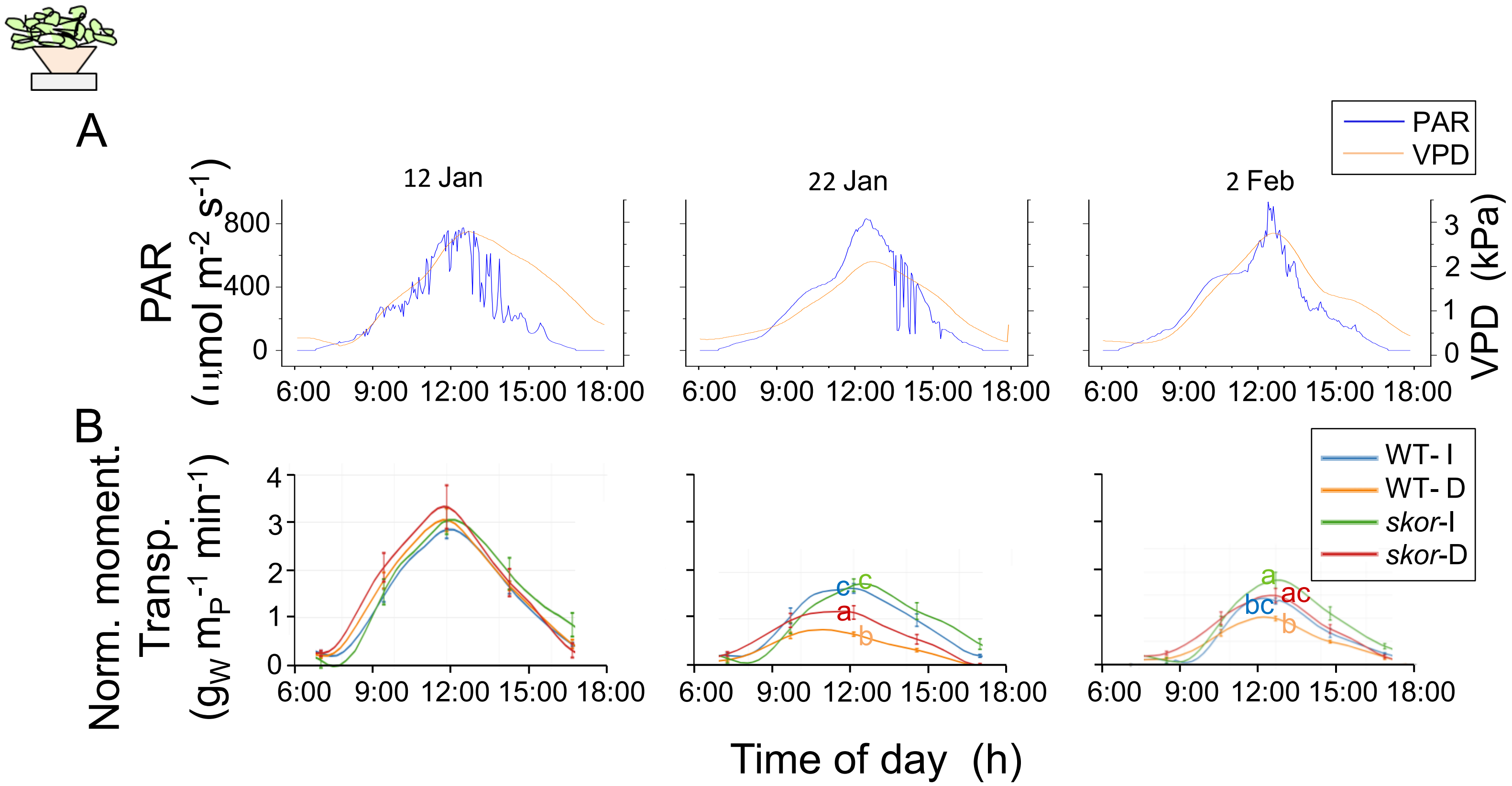


**FIGURE S7. The normalized momentary transpiration of greenhouse-grown plants on three representative days during the experiment of Fig. 5C-F** (vertical arrows in Fig. 5F indicate the dates selected based on their PAR and VPD similarities)**. A**. The atmospheric conditions during the three days. Note the wintertime hours of daylight (PAR > 0). **B.** Normalized momentary transpiration averaged over 4-7 pots holding 4 plants each, in units of (for each pot) g of lost water ($g_W$), normalized to the calculated weight, in g, of the four plants ($g_{4P}$). One group of WT plants and one group of *skor* mutant plants was water-sufficient throughout the 33-day experiment (WT-I, *skor*-I), while similar two groups were water deficient during 13 days (WT-D, *skor*-D). Jan the $12^{th}$: two days before water deprivation, Jan the $22^{nd}$: the $9^{th}$ day of water deprivation, Feb the $2^{nd}$: the $7^{th}$ day of re-irrigation. The normalized momentary transpiration of the four groups at noon (12:00) was compared on each of the three days by one-way ANOVA (Tukey HSD test) and different values (P<0.05) are indicated by different letters. Note that at noon, the *skor* mutant plants transpired more, compared to the WT, during the water stress period and even during the re-irrigation. ***Inset***: The greenhouse-grown plants (four to a pot) schematics.

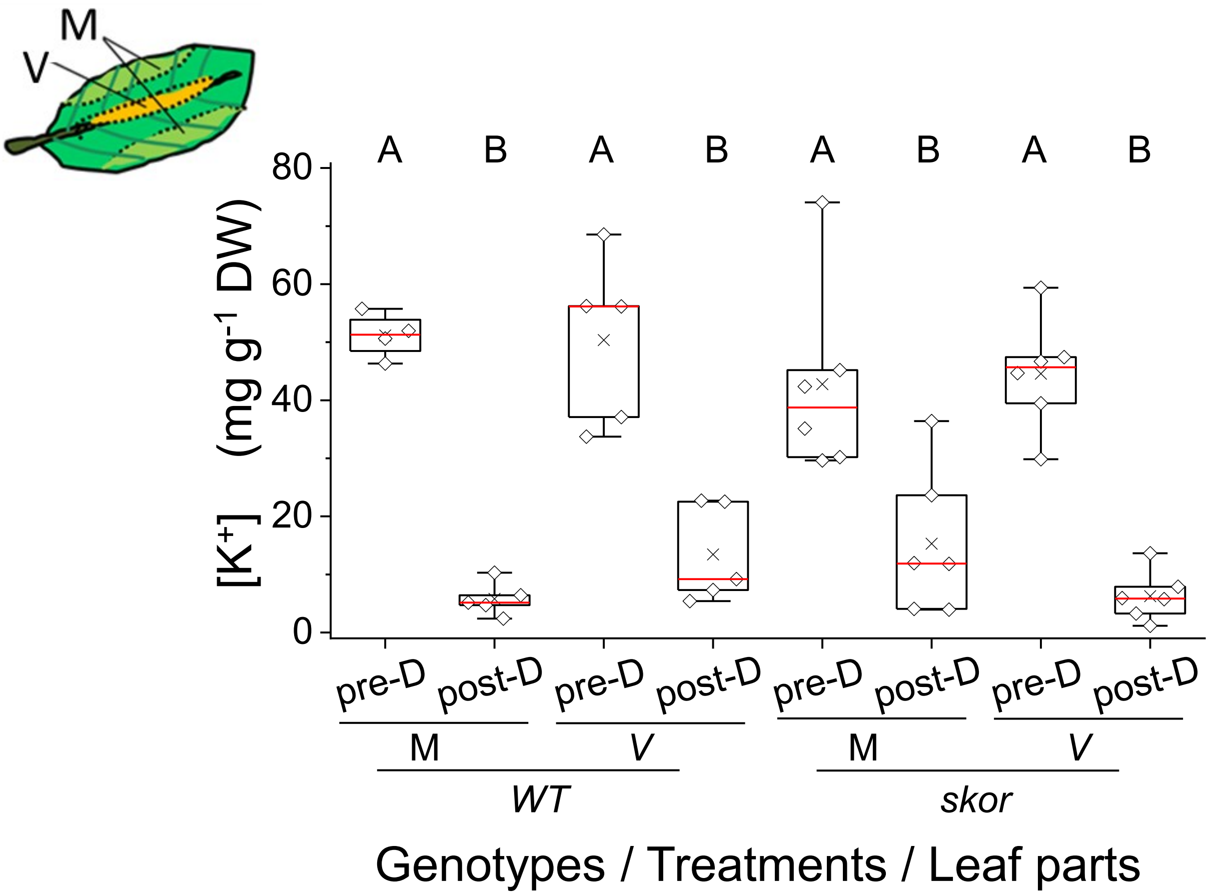


**FIGURE S8. $K^+$ accumulation in fragments of leaf blades excised from the same WT (Col) and *skor* (Col) mutant Arabidopsis plants before and after a two weeks irrigation withholding. Inset**: the excised parts definition: the edge mesophyll fragment (M) and the midrib vein fragment (V). $K^+$ concentration in leaf fragments ([$K^+$]) was determined by ICP-EOS. The pre-droughted (pre-D) plants were sampled at eight weeks of growth, during water sufficiency, and *the same* plants were sampled after two weeks of water withdrawal, at 10 weeks of growth. Two leaves per plant were considered technical repeats and their [$K^+$] values were averaged. Separate pots yielded independent biological repeats. **A.** [$K^+$] in WT (Col) leaf fragments. **B.** [$K^+$] in *skor* (Col) leaf fragments. Note the general decrease of $K^+$ content in the droughted plants leaves. **C.** *: P=0.02423 (by non-paired t-test on the ratios, *two*-tailed, equal variance), or P= 0.01242 (by a similar t-test on *log10* of the ratios), ns: not significant. Note the drought-induced *increase* of vein / mesophyll ratio in WT leaf fragments, and the absence of such an increase in *skor*. *Supports Fig. 6B.*

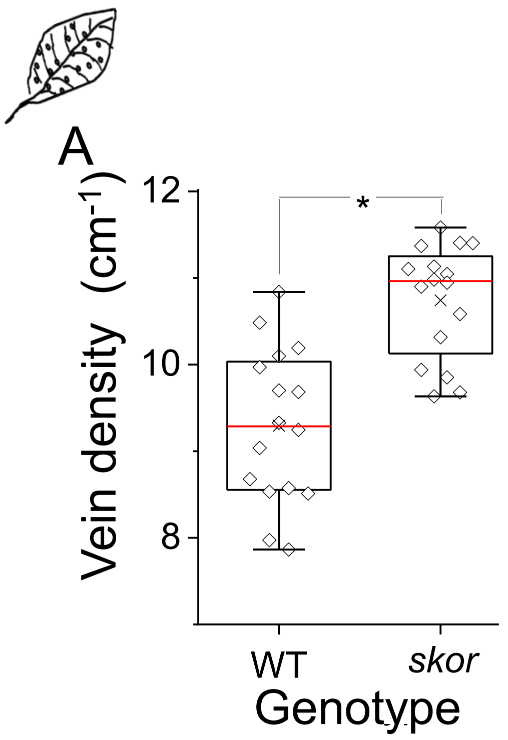


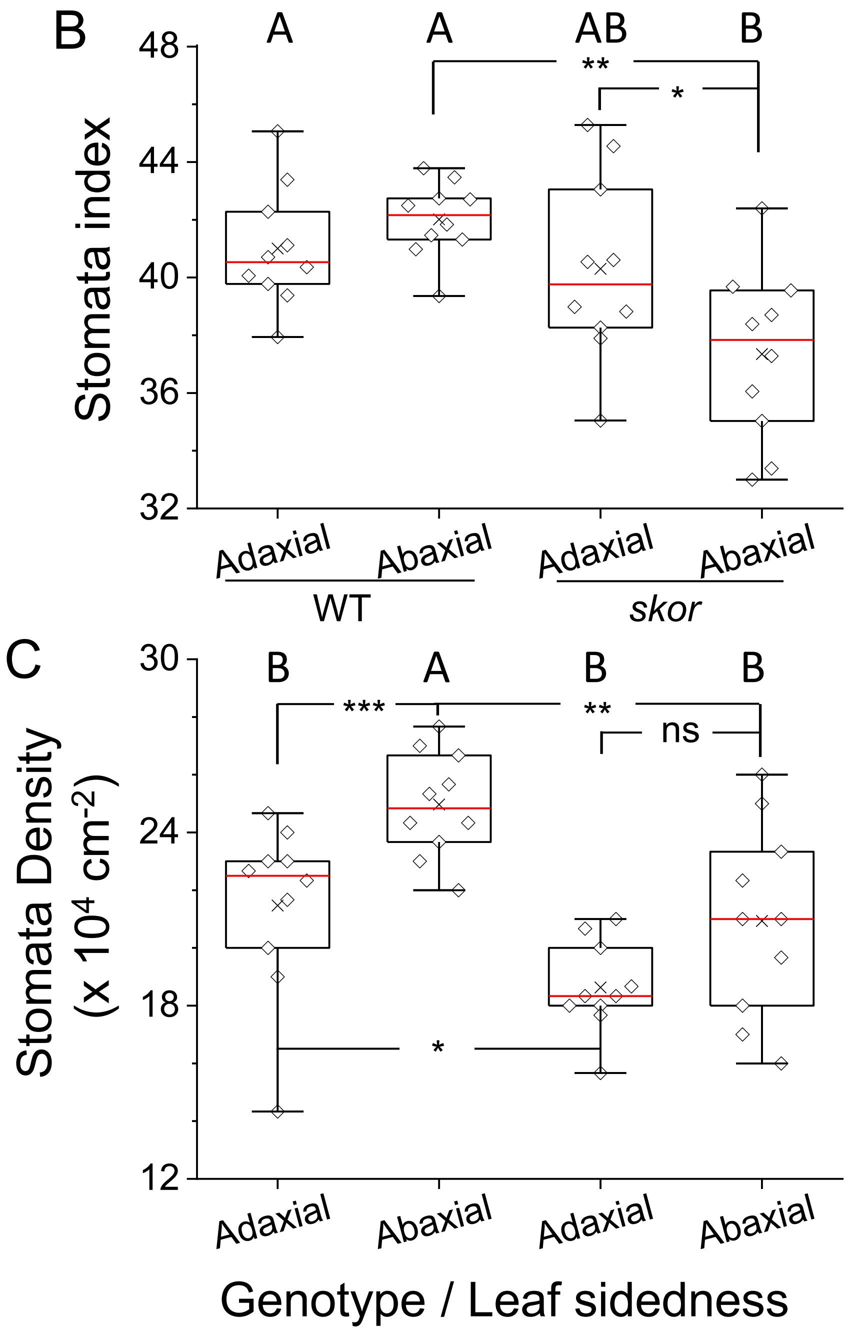


**FIGURE S9. Anatomical comparisons between leaf blades of WT (Col) and *skor* (Col). Inset.** A schematics of a leaf processed for analysis. **A.** Leaf vein density, determined for mature leaves of plants aged between 6-8 weeks using the WinRhizo software (as described by Grunwald et al., 2021). Nine plants of each WT and *skor* were sampled, 1-2 leaves from each plant, altogether 16 and 15 leaves, respectively (biological repetitions). The vein density was determined for two halves of each leaf (two technical replicates) and the values were averaged (biological repeats). *: a significant difference at P = 2.59E-06 (t-test, two-tailed, equal variance). Note the higher vein density in the leaves of *skor* (roughly 10.7 $cm^{-1}$ vs. 9.3 $cm^{-1}$ of WT).
**B.** Stomata index (the ratio of the number of stomata to the total number of stomata and epidermal cells). Different letters indicate different means (ANOVA, Tukey test, $P < 0.05$). The stomata and epidermal cells were counted on both leaf sides in triplicates (technical replicates) from leaf areas of 0.01 $mm^2$ under a 40X microscope objective and the values were averaged (biological repeats). *: P=0.046723, **: P= 0.000263, by t-test as in A.
Note the lower stomata index of the abaxial side of the *skor* leaf compared to the adaxial side of *skor* and the abaxial side of WT. **C.** Stomata density based on counts in B. *: P=0.01725, **: P= 0.00352, ***: P=0.00584, by t-test as in A. Note the 16 % lower stomata density on the abaxial leaf side of *skor* compared to the abaxial side of WT leaf. Also note that while the stomata density in WT leaf is 16 % higher on the bottom than on the top side, in *skor* the stomata density on the abaxial side might be 12% higher than on the adaxial side, but only at P=0.06403, therefore deemed nonsignificant, ns. *Supports Fig. 5B.*

Suppl. Fig. S9, Torne-Srivastava et al., 2023
