## Supplemental table for "A tale of two pumps: Blue light and ABA alter Arabidopsis leaf hydraulics via bundle sheath cells’ H^+^-pumps and channels"

### SUPPLEMENTAL MATERIALS

#### Supplemental Table S1

***The calculated free concentrations of ions in the patch-clamp pipette solutions listed in Table 1***

| **(1) pH$_{CYT}$ 7.5, nominal [Ca$^{2+}$]$_{CYT}$ 100 nM**<br>temp. 22 °C, ionic strength 0.178 N | | | **(2) pH$_{CYT}$ 7.5 , nominal [Ca$^{2+}$]$_{CYT}$ 600 nM**<br>temp. 22 °C, ionic strength 0.180 N | | |
|---|---|---|---|---|---|
| Ion | Free conc. (mM) | Total conc. (mM) | Ion | Free conc. (mM) | Total conc. (mM) |
| $Ca^{2+}$ | **1.32 e-4** | 0.7 | $Ca^{2+}$ | **5.77 e-4** | 1.4 |
| $Mg^{2+}$ | 2.0322 | 4 | $Mg^{2+}$ | 2.0683 | 4 |
| $ATP^{2-}$ | 0.1004 | 2 | $ATP^{2-}$ | 0.0993 | 2 |
| $BAPTA^{4-}$ | 1.232 | 2 | $BAPTA^{4-}$ | 0.5689 | 2 |
| **(3) pH$_{CYT}$ 8, nominal [Ca$^{2+}$]$_{CYT}$ 100 nM**<br>temp. 22 °C, ionic strength 0.179 N | | | **(4) pH$_{CYT}$ 8, nominal [Ca$^{2+}$]$_{CYT}$ 600 nM**<br>temp. 22 °C, ionic strength 0.181 N | | |
| Ion | Free conc. (mM) | Total conc. (mM) | Ion | Free conc. (mM) | Total conc. (mM) |
| $Ca^{2+}$ | **1.27 e-4** | 0.7 | $Ca^{2+}$ | **5.57 e-4** | 1.4 |
| $Mg^{2+}$ | 2.0246 | 4 | $Mg^{2+}$ | 2.0618 | 4 |
| $ATP^{2-}$ | 0.0948 | 2 | $ATP^{2-}$ | 0.0937 | 2 |
| $BAPTA^{4-}$ | 1.230 | 2 | $BAPTA^{4-}$ | 0.5679 | 2 |

The values have been calculated using the free program Extended MaxChelator https://somapp.ucdmc.ucdavis.edu/pharmacology/bers/maxchelator/webmaxc/webmaxcE.htm

The ionic strength was calculated with the following additional information on the speciation of MES and HEPES (from WebBufferCalc v1.00, https://somapp.ucdmc.ucdavis.edu/pharmacology/bers/maxchelator/webmaxc/webbufcalc.htm for bath solutions with MES (pKa=6.16 at 20 °C) at pH 5.6, 39 % of MES was in the -1 form, while at pH ≥7, 94% of MES was in the -1 form, and for solutions with HEPES (pKa=7.55 at 20 °C), at pH 7.5, 68 % of HEPES was in the -1 form and at pH 8, 87 % of HEPES was in the -1 form; Bers et al., 2010).

**Bers DM, Patton CW, Nuccitelli R.** (2010) A practical guide to the preparation of $Ca^{2+}$ buffers. Methods Cell Biol. 99:1-26.
